# Modulation of SLFN11 induces changes in DNA Damage response

**DOI:** 10.1101/2023.04.02.535254

**Authors:** Christophe Michel Raynaud, Eiman I. Ahmed, Ayesha Jabeen, Apryl Sanchez, Shimaa Sherif, Tatiana Carneiro Lobo, Amany Awad, Dina Awartani, Adviti Naik, Remy Thomas, Julie Decock, Gabriele Zoppoli, Davide Bedongnetti, Wouter Hendrickx

**Affiliations:** Pediatric Cancer Omics Lab, Cancer group, Research Branch, Sidra Medicine, Doha, Qatar; Cancer Immunogenetics Lab, Cancer group, Research Branch, Sidra Medicine, Doha, Qatar; Clinical and Experimental Oncology and Hematology, Ospedale Policlinico San Martino, Genova, Italy; Translational Cancer and Immunity Center, Qatar Biomedical Research Center, Doha, Qatar; College of Health and Life Sciences (CHLS), Hamad bin Khalifa University (HBKU), Doha, Qatar; Department of Internal Medicine (DiMI), University of Genoa and Ospedale Policlinico San Martino, Genoa, Italy; Ospedale Policlinico San Martino IRCCS per l'Oncologia, Genoa, Italy

## Abstract

**Background:** Lack of Schlafen family member 11 (SLFN11) expression has been recently identified as a dominant genomic determinant of response to DNA damaging agents in numerous cancer types. Thus, strategies aimed at increasing SLFN11 could be used to restore chemosensitivity of refractory cancers.

As oncogenic downregulation is often driven by methylation of the promotor region, we explore the demethylation effect of 5-aza-2’-deoxycytidine (decitabine), on the SLFN11 gene methylation. Since SLFN11 has been reported as an interferon inducible gene, and interferon is secreted during an active anti-tumor immune response, we investigated the in vitro effect of IFN-γ on SLFN11 expression in breast cancer cell lines. A second broader approach to show cross talk between immune cells and SLFN11 expression is indirect co-culture of breast cancer cells with activated PBMCs and evaluate if this can drive SLFN11 upregulation. Finally, as a definitive and specific way to modulate SLFN11 expression we implemented SLFN11 dCas9 (dead CRISPR associated protein 9) systems to specifically increase or decrease SLFN11 expression.

**Results:** We first confirmed a correlation previously reported between methylation of SLFN11 promoter and its expression across multiple cell lines. We showed *in-vitro* that decitabine and IFN-γ could increase moderately the expression of SLFN11 in both BT- 549 and T47D cell lines, but not in strongly methylated cell lines such as MDA-MB-231. Though, *in-vitro*, the co-culture of the same cell lines with CD8-CD25 activated PBMC failed to increase SLFN11 expression. On the one hand, the use of a CRISPR-dCas9 UNISAM system could increase SLFN11 expression significantly (up to 5-fold), stably and specifically in BT-549 and T47D cancer cell lines. Though, this system also failed to force a strong expression of SLFN11 in cell lines with robust SLFN11 promoter methylation such as MDA-MB-231. On the other hand, the use of CRISPR-dCas9 KRAB could significantly reduce the expression of SLFN11 in BT-549 and T47D. We then used the modified cell lines to confirm the alteration in chemo sensitivity of those cells to treatment with DNA Damaging Agents (DDAs) such as Cisplatin and Epirubicin or DNA Damage Response (DDRs) drugs like Olaparib. RNAseq was used to elucidate the mechanisms of action affected by the alteration in SLFN11 expression.

**Conclusion:** To our knowledge this is the first report of the stable non-lethal increase of SLFN11 expression in a cancer cell line. Our results show that induction of SLFN11 expression can enhance DDA and DDR sensitivity in breast cancer cells and dCas9 systems may represent a novel approach to increase SLFN11 and achieve higher sensitivity to chemotherapeutic agents, improving outcome or decreasing required drug concentrations. SLFN11-targeting therapies might be explored pre-clinically to develop personalized approaches.

## Introduction

Schlafen 11 (SLFN11), is a highly conserved mammalian nuclear protein found to be an essential component during replication stress. In brief, SLFN11 induces an irreversible replication block and eventually apoptosis (Mu et al. 2016). SLFN11 binds to Replication Protein A (RPA) in the stalled replication forks. It interacts with MCM3 (Minichromosomal maintenance complex component 3), and DHX9 (DExH-box helicase) (Murai et al. 2018; Nogales et al. 2015). This leads to the opening of the chromatin around the replication initiation sites, which activates the transcription of immediate early genes that can induce cell cycle arrest. Thereby blocking any further replications from occurring (Murai et al. 2020) This is done independently of, and in parallel with, the ATR-CHEK 1 S-phase checkpoint in the DDR pathway (Murai et al. 2018).

SLFN11 expression has been strongly correlated with sensitivity to DNA damaging agents (DDA), like platinum salts such as cisplatin, or agents affecting DNA Damage Repair (DDR) like PARP inhibitors such as Olaparib and to topoisomerase I and II inhibitors such as topotecan and Epirubicin (Barretina et al. 2012; Zoppoli et al. 2012). More recently SLFN11 immunohistochemistry of ovarian cancer tissue was able to predict response to platinum-based chemotherapies (Winkler et al. 2021). Higher SLFN11 gene expression showed a better prognosis in breast cancer (Isnaldi et al. 2019). Hence, SLFN11 could make for a good predictive biomarker of therapeutic response or a treatment target in many cancers including breast cancer.

Next to a predictive biomarker SLFN11 could be a therapeutic target for sensitizing cancer cell to chemo, radio, or immunotherapy. To comprehensively quantify the effect of the gene’s re-induction on the cells chemosensitivity to different drugs we modulate the SLFN11 expression in several ways.

The SLFN11 gene expression can be suppressed by three epigenetic mechanisms: promotor methylation (Nogales et al. 2015), histone deacetylation (Tang et al. 2018), and histone methylation by the Polycomb Repressive Complex (PRC) (Stewart et al. 2017). Most cell lines that lack SLFN11 expression were found to feature hypermethylation-associated silencing in the CpG island in the promotor region of the SLFN11 gene. This silencing correlates with increased resistance to platinum chemotherapies drugs (Nogales et al. 2015). This prompted us to attempt demethylation of the promotor region of SLFN1 using decitabine.

In addition, SLFN11 has been identified as an interferon (IFN)-stimulated gene (Mavrommatis, Fish, and Platanias 2013). SLFN11 gene expression is induced upon interferon signaling in case of viral infections such as HIV or Zika virus (Li et al. 2012). Therefore, we tried inducing SLFN11 expression using in-vitro Interferon Gamma stimulation in our breast cancer cells.

This paper comprehensively investigates the relationship between modulation of SLFN11 expression either by interferon, decitabine demethylation, co-culture with activated PBMC’s or CRISPR alteration and the resulting changes in chemosensitivity. Hypothesizing that in cancers which are resistant to chemotherapy, upregulation of SLFN11 can restore their chemosensitivity. Once cell lines were selected of the multiple methods investigated to upregulate SLFN11 expression, the most robust modulation was achieved with CRISPR, which was precise, effective, and lacked wider transcriptomic effects.

## Material and methods

### Cell lines and culture

All cell lines were purchased from ATCC. (**Table 1**) All cell lines were culture in advanced RPMI (Gibco, #12633012) complemented with 10% FBS (Sigma, #), Glutamax (Gibco, #35050061) and antibiotic-antimycotic (Gibco, #15240096). Cells were cultured at 37°C, 5% CO2 and 95% humidity. Cells were detached using TrypLE express enzyme (Gibco, #12605036). Human foreskin fibroblasts were used across the manuscript as positive control and reference for expression of SLFN11.

**Table 1:**
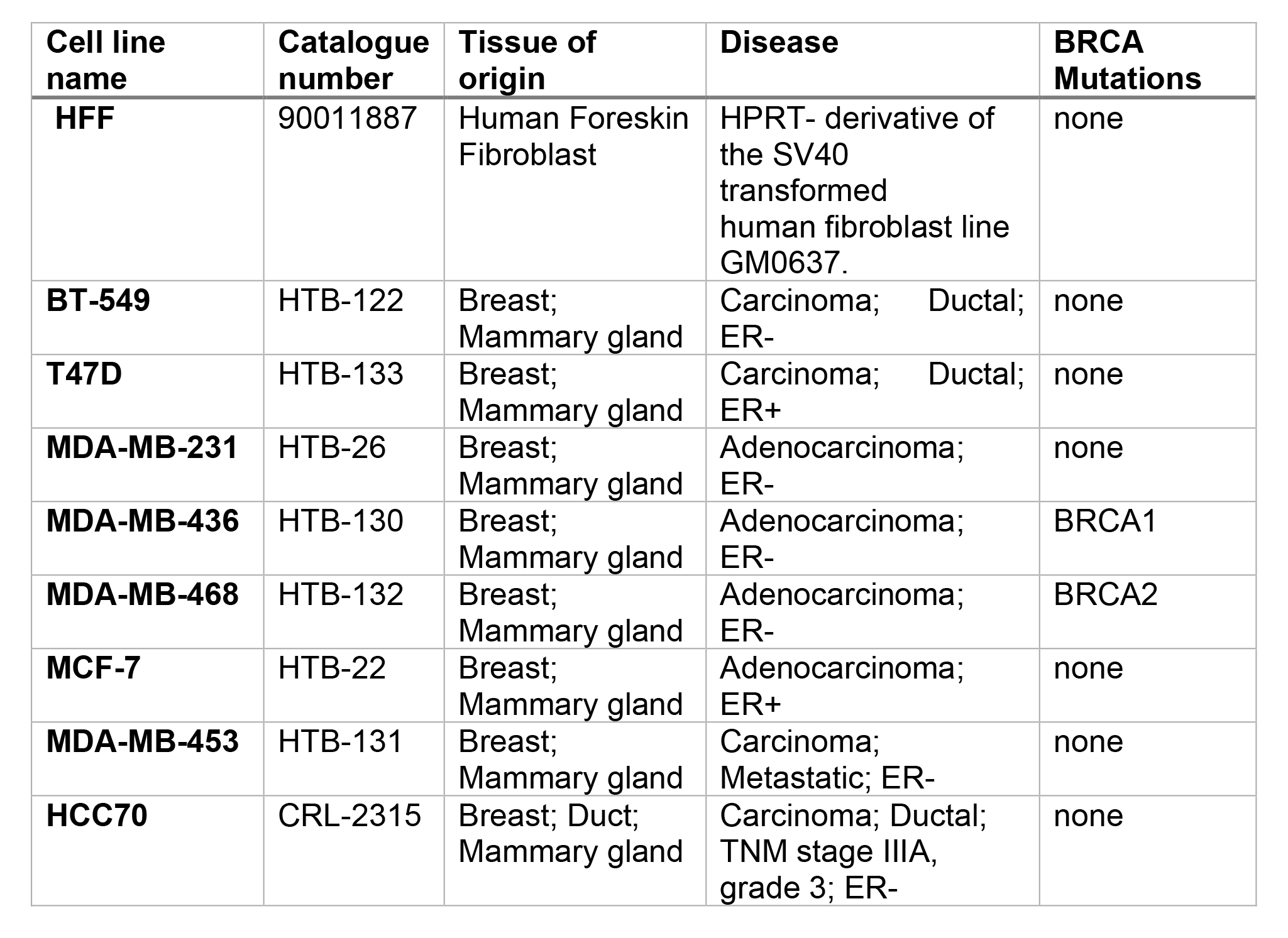
cell lines

### DNA/RNA extraction

### Q-PCR

1ug of RNA was used for reverse transcription using TaqMan™ Reverse Transcription Reagents (Invitrogen, #N8080234) and random hexamers. cDNA was diluted 20 times with DNA/RNA free water. TaqMan™ Gene Expression Master Mix (Applied bioscience, #4369016) and Hs03003631_g1 (for Eukaryotic 18S rRNA) and Hs00536981_m1 (for SLFN11) TaqMan probes (Thermo scientific, #4331182) were used according to manufacturer recommendation. Real time PCR was run in 96 well plates on QuantStudio 12K flex system (Thermofisher Scientific). Q-PCR was done in triplicate for each sample and data were analyzed by gene expression comparison using ΔΔCT on (QuantStudio 12K Flex Realtime PCR system V1.2.2) using S18 as the housekeeping gene.

### Western blot

After culture, 5 × 10^6^ cells of cells were washed with DPBS and lysed with 400ul of RIPA Lysis and Extraction Buffer (Thermo Scientific, #89900) complemented with Halt™ Protease Inhibitor Cocktail (Thermo scientific, #78430) and sonication for 30 sec. cell debris was removed by 30 min centrifugation at 14.000g. Supernatant containing protein extract were kept at −20°C until use. Protein concentration was assessed using Pierce BCA protein assay kit (Thermo scientific, #23225). Capillary western blot was done using a Wes system (protein simple) with 12-230 kDa Separation Module, 8 x 25 capillary cartridges (Protein simple, #SW-W004), EZ Standard Pack 2 (Protein simple, #PS- ST02EZ-8) and Anti-mouse detection module (Protein simple DM-002). Mouse anti human SLFN11 (Santa Cruz, #SC-515071) and anti β-actin (Licor, #926-42212) both diluted at 1 in 100 were used as primary antibody.

Analysis was done using compass for Simple western (ProteinSimple, V5.0.0) and area of histogram peaks were used for quantification. All western blot analysis were normalized for β-actin expression.

### Promoter Methylation Analysis by MSP

Promoter methylation was analyzed using Methyl Specific Polymerase Chain Reaction (MSP). Genomic DNA was extracted as previously described, bisulfite conversion was performed using EZ DNA methylation kit (Zymo research, # D5001). PCR was performed using the primers: Forward Methylated specific primer (GTAGCGGGGTAGAAAAGTAGAAC) and Reverse Methylated specific primer (TAAAATTTAACGACGACCGATACG) for methylated specific PCR with a PCR product of 108bp. Forward Unmethylated specific primer (GTAGTGGGGTAGAAAAGTAGAAT) and Reverse unmethylated specific primer (TAAAATTTAACAACAACCAATACA) for unmethylated specific PCR with a PCR product of 105bp, 1ul of converted DNA and AmpliTaq Gold™ 360 Master Mix (Applied Biosystems, #4398876).

The PCR product was then run on 2% agarose (Sigma, #A4718) gel containing Ethidium Bromide (Sigma, #E1510) and picture were taken using Chemidoc XRS system (Biorad, # 1708265) with the single channel ethidium bromide agarose gel protocol. Band intensity was measured using Image J (https://imagej.nih.gov/ij/, 1997-2018. Schneider, C.A., Rasband, W.S., Eliceiri, K.W. "NIH Image to ImageJ: 25 years of image analysis") and relative intensity between methylated and unmethylated specific PCR was calculated.

### Drugs

5-Azacytidine (Decitabine) (DAC) (MP Biomedicals, #154803) was reconstituted at 20mM in DMSO (Sigma, #D4540) and used at 5uM final concentration; IFN-γ (Peprotech, #300- 02-100UG) was diluted in water at 10μM and used at 5nM final concentration, cis-Diamineplatinium (II) dichloride (Cisplatin) (Sigma, #479306) was reconstituted fresh at 2mM in NaCl solution 0.9% (Sigma, #SW8776) and used at various concentration as indicated; Epirubicin (Sigma,# 1237382) was reconstituted at 3.5mM in water and used at various concentration; Olaparib (Biovision, #1952-25) was reconstituted at 57.54mM in DMSO and used at various concentration.

### Indirect co-culture model

A total of 5 × 10^4^ cancer cells (BT-549, T47D and MDA-MB-231) were seeded per well in a 24-well plate and cultured overnight. Total PBMCs were plated in a flat-bottom multi-well plate (Thermo Fisher Scientific, Nunclon Δ Surface) and incubated for 2h at 37°C and 5% CO_2_ and following the incubation, the non-adherent peripheral blood lymphocyte population (PBLs) were isolated. Next, the non-adherent PBLs were activated overnight using 2 μg/ml of plate-bound anti-human CD3 and CD28 antibodies (eBioscience) at 37°C and 5% CO_2_. To set up the indirect co-culture, the activated PBLs were placed on top using transwell inserts with 0.4 µm pore size (Corning) and a Target:Effector (T:E) ratio of 1:20 to enable exchange of soluble factors between cancer cells and PBLs without direct cell-cell contact. Cancer cells were plated alone without PBls as control. The cells were co-cultured for 72 h at 37°C and 5% CO2, after which RNA was isolated from cancer cells for downstream analysis.

### CRISPR cell engineering - gRNA design

IDT custom gRNA design tool was used to design gRNA along the core region of the promoter, distributed along the promoter as illustrated in Supplementary figure 1.

The sequence of the gRNA is as follow:

**Table 2:**
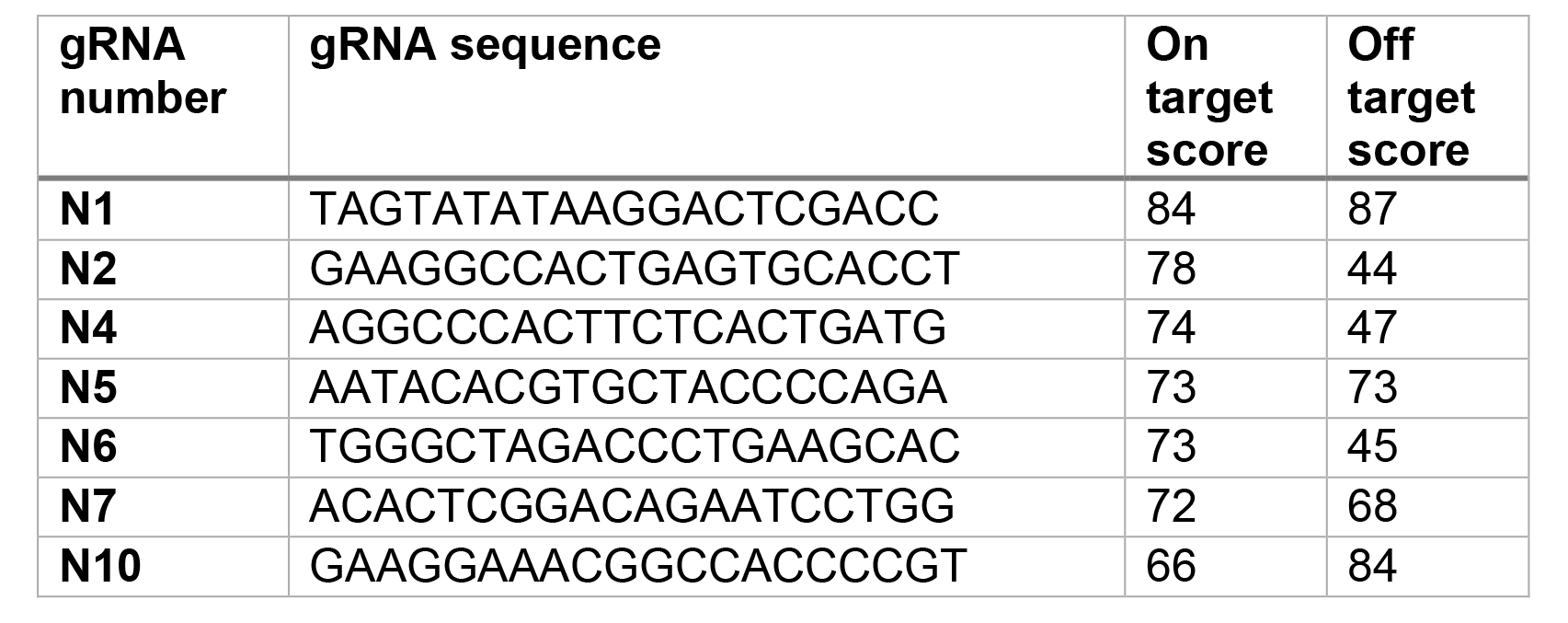
gRNA sequence

The sequence of the scramble (SCR) gRNA used is GCACTACCAGAGCTAACTCA.

Forward and reverse primers were ordered accordingly to be inserted in the appropriate plasmids as described below.

### CRISPR cell engineering - Cloning of gRNA into UNISAM and KRAB plasmids

PB-UniSAM containing mCherry was a gift from Lesley Forrester (Addgene plasmid # 99866; http://n2t.net/addgene:99866; RRID: Addgene_99866) (Fidanza et al. 2017). pLV hU6-sgRNA hUbC-dCas9-KRAB-T2a-GFP was a gift from Charles Gersbach (Addgene plasmid # 71237; http://n2t.net/addgene:71237; RRID: Addgene_71237) (Thakore et al. 2015). Cloning in PB-UniSAM was done using BbsI (New England Biolabs, #R3539L) as indicated in (Fidanza et al. 2017). Cloning into pLV hU6-sgRNA hUbC-dCas9-KRAB-T2a- GFP was performed using BsmBI restriction enzyme (New England Biolabs, #R0580L) as indicated in (Thakore et al. 2015). Briefly, gRNAs were obtained from Integrated DNA technology as two single strand oligos and annealed in 40 μl of annealing buffer (Origene, #GE100007) with 2 μl of each oligo (100 μM) and annealed in the thermal cycler at 95 °C for 4 minutes followed by cooling to 25 °C with 1°C/minute ramp. Annealed oligos were then diluted 10 times in water. 10ng of purified *BbsI* linearized UniSAM or BsmBI linearized pLV hU6-sgRNA hUbC-dCas9-KRAB-T2a-GFP was ligated with 1 μl of diluted annealed gRNA with 0.5 μl of T4 ligase (New England Biolabs, #M0202L) in a total volume of 10 μl. Solution was incubated at room temperature for two hours prior to transformation in E.Coli STBL3 (Invitrogen, # C737303). Minipreps were performed using QIAprep Spin Miniprep Kit (Qiagen, #27106X4). For stable modification of cells using UniSAM, co-transfection was performed with pcDNA3-transposase gifted by Dr. Juan Cadinanos. Correctly assembled UniSAM and pLV hU6-sgRNA hUbC-dCas9-KRAB- T2a-GFP vectors were confirmed by complete Sanger sequencing using BigDye™ Terminator v3.1 Cycle Sequencing Kit (Applied Biosystem, #4337455) and DyeEx 2.0 Spin kit (Qiagen, #63204) according to manufacturer recommendation and analyzed on ABI3500xL (Applied biosystem, #4406016).

### CRISPR cell engineering - Cell transfection

Electroporation was performed using Neon transfection system (Invitrogen, #MPK5000) with Neon™ Transfection System 10 µL Kit (Invitrogen, #MPK1096) using 2ug of DNA for 1.10^5^ cells in 24 well plate. Electroporation protocol for each cell line was identified using pmaxCloningTM vector (Lonza, # VDC-1040). The optimal protocol for each cell line is indicated here below:

**Table 3:** electroporation conditions for each cell line

| Cell line | Voltage | Width | Pulses |
| --- | --- | --- | --- |
| **BT-549** | 1300 | 30 | 1 |
| **T47D** | 1400 | 30 | 1 |
| **MDA-MB-231** | 1600 | 20 | 1 |

After 1 week of culture, cells were further purified by sort.

### CRISPR cell engineering - Sort of cells

Cells were harvested and blocked in PBS with 5%FBS and 1%BSA and cell clumps removed on 40uM cell strainer (Falcon, #382235). Single-cell suspension was analyzed and sorted on SORP FACSAriaIII (BD Biosciences Special Order Research Product). Data were processed with BD FACSDiva™ Software V8.0.1 (BD Biosciences). GFP fluorescence was acquired with 488 nm blue laser and 530/30 nm emission filter, and mCherry fluorescence was acquired with 561 nm yellow-green laser and 610/20 nm emission filter. During cell-sorting 4-way purity-phase mask was applied. To ensure maximum purity, cells were serially sorted 3 times prior analysis and use.

### Viability analysis

Cells were grown in 96 well plate for 24 to 72 hours accordingly with or without treatment. 3000 and 5000 cells were plated for BT-549 and T47D respectively 24h prior treatment. Viability was assessed using ATPlite Luminescence Assay System (Perkinelmer, #6016949) and Calcein AM (Invitrogen, #C3099). Luminescence was measured with Ensight plate reader (Perkinelmer, #HH34000000). Calcein AM was assessed by fluorescence intensity measurement by well scan from bottom with excitation at 494 nm and emission at 517 nm on Ensight plate reader (Perkinelmer, #HH34000000).

### mRNA sequencing

mRNA-sequencing was performed using QuantSeq 3’ mRNA-Seq Library Prep Kit FWD for Illumina (Cat. 015.96) (75 single-end) with a read depth of average 9.31 M, and average read alignment of 55.8%. Single samples were sequenced across multiple lanes, and the resulting FASTQ files were merged by sample. All samples passed FastQC (v. 0.11.8) were aligned to the reference genome GRChg38 using STAR (v. **2.6.1d**) (Dobin et al. 2013). BAM files were converted to a raw counts expression matrix using HTSeq-count (v. 0.9.1) (Anders, Pyl, and Huber 2015)

### RNAseq Data Normalization

Normalization was done using R Bioconductor package EDAseq (Exploratory Data Analysis and Normalization for RNA-Seq) (v. 2.28.0) (Risso et al. 2011) to remove within and between lane effects. Data was then quantile normalized using R Bioconductor package preprocessCore package (v. 1.56.0) (Bolstad 2016) and log2 transformed. All downstream analysis was done using R (v. 4.1.2). Batch effects were removed for each cell line separately using ComBat () function from R Bioconductor package sva (v. 3.42.0) (Leek et al. 2012). Genes with row sum equal to zero were removed before applying ComBat. Data were then combined, and quantile normalized again as described previously. Principal component analysis was done based on genes expression to assess global transcriptional differences between the samples using prcomp() function and plotted using R CRAN package ggplot2 (v. 3.3.5) (Wickham 2009).

### Differentially expressed genes

Differentially expressed genes analysis was performed on log2 normalized mRNA expression data using R Bioconductor package limma (v. 3.50.0) (Ritchie et al. 2015) with Benjamini-Hochberg (B-H) FDR. Within each comparative analysis, genes with row sum equal to zero were removed. To visualize the overlap of differentially expressed genes between the conditions, R CRAN package VennDiagram (v. 1.7.1) was used (Chen and Boutros 2011). Differentially expressed genes were then plotted in a heatmap using R Bioconductor package ComplexHeatmap (v. 2.10.0) (Gu et al. 2014).

### Single Sample Gene Set Enrichment Analysis (ssGSEA)

ssGSEA was done using normalized, log2 transformed data for the selected list of genes. Enrichment score (ES) was calculated using gsva function from R Bioconductor package GSVA (v. 1.42.0) (Hänzelmann, Castelo, and Guinney 2013). ES was calculated for genes obtained from DEG analysis using Limma. Gene sets to reflect enrichment of apoptosis and glycogen metabolism pathways were downloaded from Molecular Signatures Database (MSigDB) (Liberzon et al. 2011).

## Results

### Baseline SLFN11 expression and associated methylation profiles across a panel of 8 different breast cancer cell lines

In order to determine SLFN11 baseline expression in different breast cancer cell lines, BT-549, T47D, MDA-MB-231, MDA-MB-436, MDA-MB-468, MCF-7, MDA-MB-453 and,HCC70 were screened for SLFN11 expression both at mRNA and protein level, an immortalized human fibroblasts cell line (HFF) was used as a normal control. Analysis by RT-qPCR confirms differential SLFN11 mRNA expression among multiple breast cancer cell lines; however, MDA-MB-231, MDA-MB-543 and HCC70 showed almost null SLFN11 mRNA expression (**Figure 1A**). To validate SLFN11 protein expression, capillary western (Wes) was conducted (**Figure 1B, Supplementary Figure 3A**). In addition, figure 1C shows a very significant correlation between mRNA and protein expression of SLFN11 analyzed by RT-qPCR and WES; (R^2^=0.86 and p= 0.0008) (**Figure 1C**).

**Figure 1:**
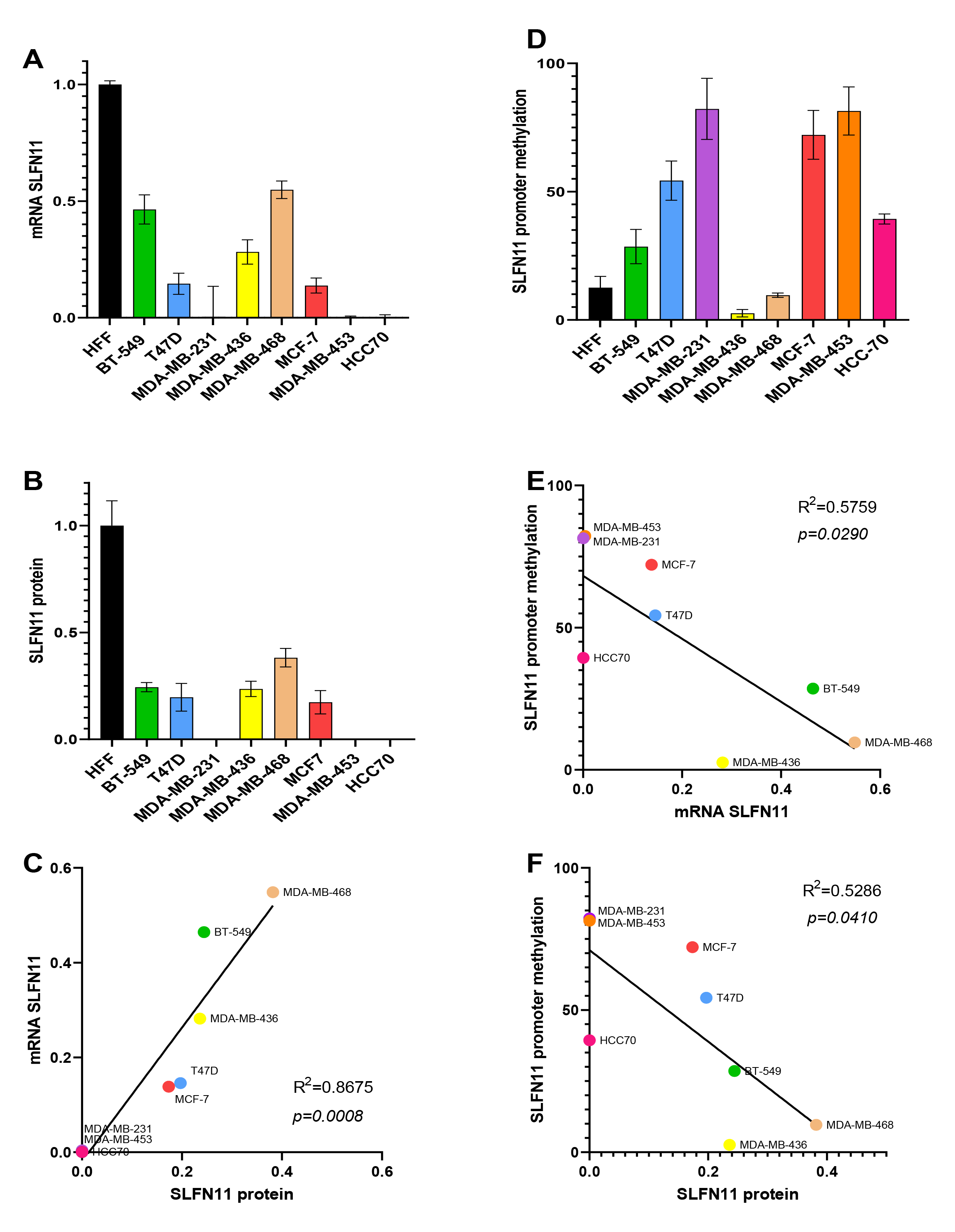
*Baseline expression of SLFN11 across a panel of 8 breast cancer cell lines and associated methylation profile* (A) Relative mRNA expression of SLFN11 analyzed by RT-qPCR. SLFN11 expression in the different cell lines shown relative to the expression level in HFF (human foreskin fibroblast) used as control cells (N=3). (B) Capillary Western immunoassay (WES) of SLFN11 expression in those breast cancer cell lines. Quantification of band intensity of SLFN11 relative to the expression level in HFF cells (N=2). (C) Correlation between SLFN11 mRNA analyzed by Q-PCR and protein expression analyzed by WES. (D) Percentage of methylation of SLFN11 promoter analyzed by methyl specific PCR (MSP) within the CPG island of the promoter (N=4). (E). Correlation of SLFN11 promoter methylation and SLFN11 mRNA expression analyzed by Q-PCR. (F). Correlation of SLFN11 promoter methylation and SLFN11 protein expression analyzed by WES.

Next, to understand what regulates SLFN11 expression, methylation analysis of SLFN11 promoter was conducted using methylation specific PCR (MSP) (**Figure 1D**), a good correlation between promoter methylation and both the mRNA and protein expression was observed (R^2^=0.57 and p=0.029; R^2^=0.53 and p=0.041) (**Figure 1E and 1F**). The data shows an increase in SLFN11 methylation, resulting in downregulated mRNA and protein expression of SLFN11,

Therefore, confirming that the methylation of SLFN11 promoter plays a vital role in regulating SLFN11 expression.

### Limited increase in SLFN11 expression upon treatment with Decitabine (DAC)

Based on the screening results we selected 3 representative breast cancer cell lines (BT- 549, T47D and MDA-MB-231) for further experiments as they cover the range of SLFN11 expression observed at baseline (null, low and moderate compared to HFF). In order to re-establish normal SLFN11 expression through demethylation of the SLFN11 gene in the selected breast cancer cell lines, we treated them with 5µm of DAC for 72h. We then analyzed SLFN11 mRNA expression using RT-qPCR which revealed limited but significant increases in expression in both BT-549 and T47D compared to DMSO treated cells. (**Figure 2A**). Though, despite a moderate increase the protein expression of SLFN11 in those cells lines, it was not significant relative to DMSO treated cells (p=0.1373 for BT-549 and p=0,284 for T47D) (**Figure 2B, Supplementary Figure 3B**). Besides, DAC treatment did not significantly affect the methylation of SLFN11 promoter in BT-549 and T47D breast cancer cell lines. In contrast, the heavily methylated MDA-MB-231 showed almost no effect of DAC on SLFN11 mRNA and protein expression. Even though, there was a significant decrease in SLFN11 promoter methylation in DAC treated MDA-MB-231 cells (p=0.0122) (**Figure 2C**).

**Figure 2:**
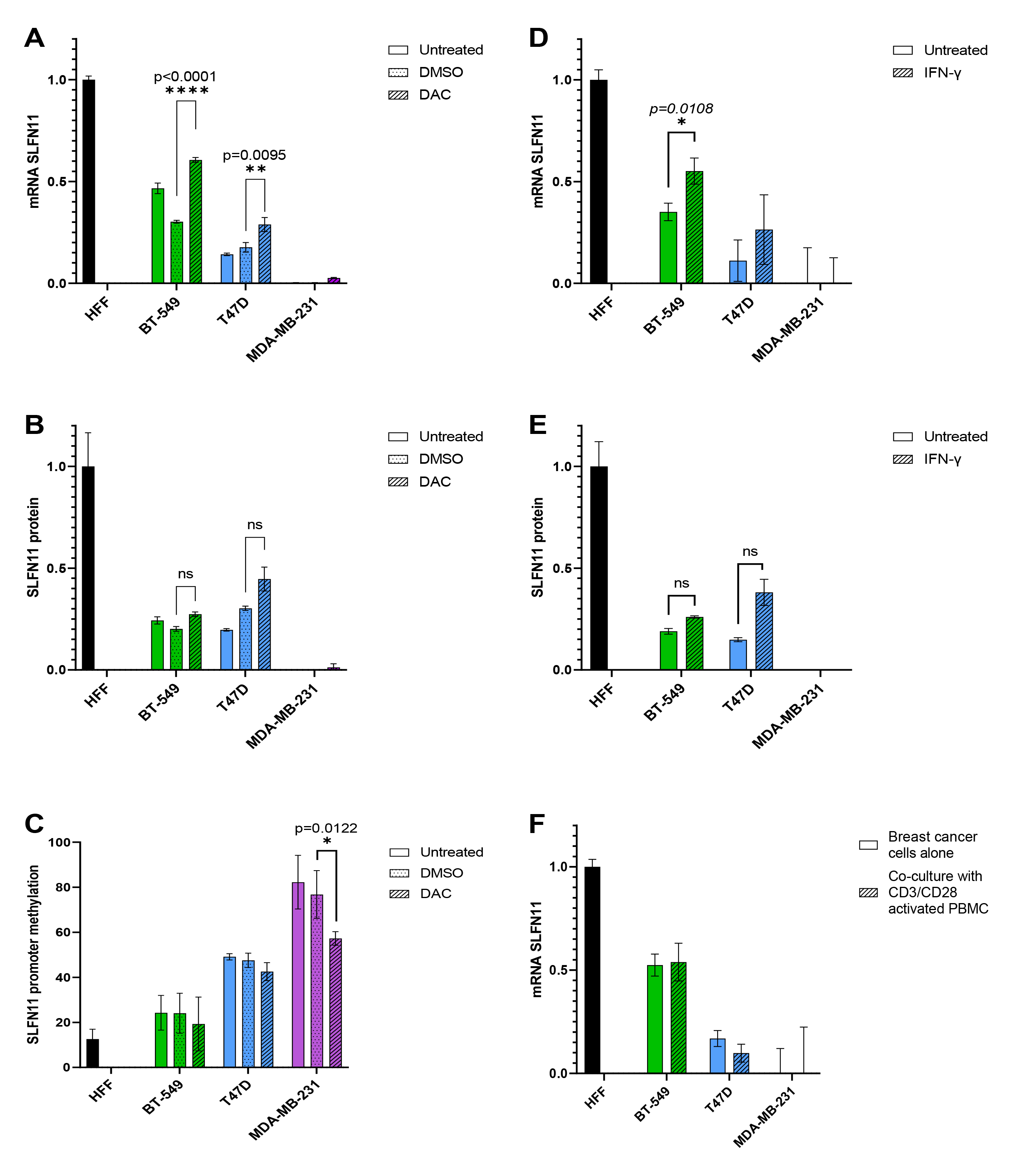
*IFN-γ and DAC have limited effect of SLFN11 expression* (A-B). Relative expression of SLFN11 analyzed by Q-PCR (N=3) (A) and by WES (N=2) (B) after treatment with 5uM of DAC for 72h compared to the expression level in untreated HFF. (C) Percentage of methylation of SLFN11 promoter analyzed by MSP after treatment with 5uM of DAC for 72h (N=4) compared to untreated HFF. (D-E) Relative expression of SLFN11 analyzed by Q-PCR (N=3) (D) and by WES (N=2) (E) after treatment with 5nM of IFN-γ for 24h compared to the expression level in untreated HFF. (F). Relative mRNA expression of SLFN11 analyzed by RT-qPCR after co-culture with CD3/CD28 for 24h compared to the expression level in HFF.

### Limited increase in SLFN11 expression upon treatment with IFN- Ɣ

Since SLFN11 expression is interferon inducible (Mezzadra et al. 2019), our next approach was to treat breast cancer cells with 5nM of IFN-Ɣ for 24h. Our data shows IFN- Ɣ could increase mRNA expression of SLFN11 in both BT-549 and T47D. However, significance (p=0.0108) could only be demonstrated in BT-549 cancer cells (**Figure 2D**). In contrast, expression level increases of SLFN11 protein were detected but non-significant in both BT-549 and T47D (**Figure 2E, Supplementary Figure 3C**).

### No increase in SLFN11 expression co-culture with activated PBMC

Finally, we attempted to induce increased SLFN11 expression by co-culturing breast cancer cells with PBMC activated by CD3/CD28 for 24h. Surprisingly, the results revealed no increment in SLFN11 mRNA expression (**Figure 2F**).

### CRISPR-dCas9 can significantly alter SLFN11 expression in BT-549 and T47D cancer cells

Since the increases in SLFN11 expression using DAC or IFN-Ɣ resulted in only minor increases we attempted modulation of SLFN11 expression using UNISAM (Unique Synergistic Activation Mediator) for dead Cas9 (dCas9) activation of SLFN11 (Fidanza et al. 2017) and KRAB (Kruppel associated box) for dCas9 inhibition of SLFN11(Parsi et al. 2017). We designed 7 gRNAs across the central region of the promoter (**Supplementary figure 1**), we then established stable expressing cell lines for all 7 gRNAs. All modified cell lines were screened by WES and Q-PCR to identify the most efficient single gRNA that could successfully modulate SLFN11 expression levels in the selected cancer cells. The gRNA “N7” was found to be most effective for upregulation of SLFN11 and therefore further used as gRNA for activation and inhibition **(Supplementary figure 2A-I, Supplementary figure 3D-G).**

**Figure 3:**
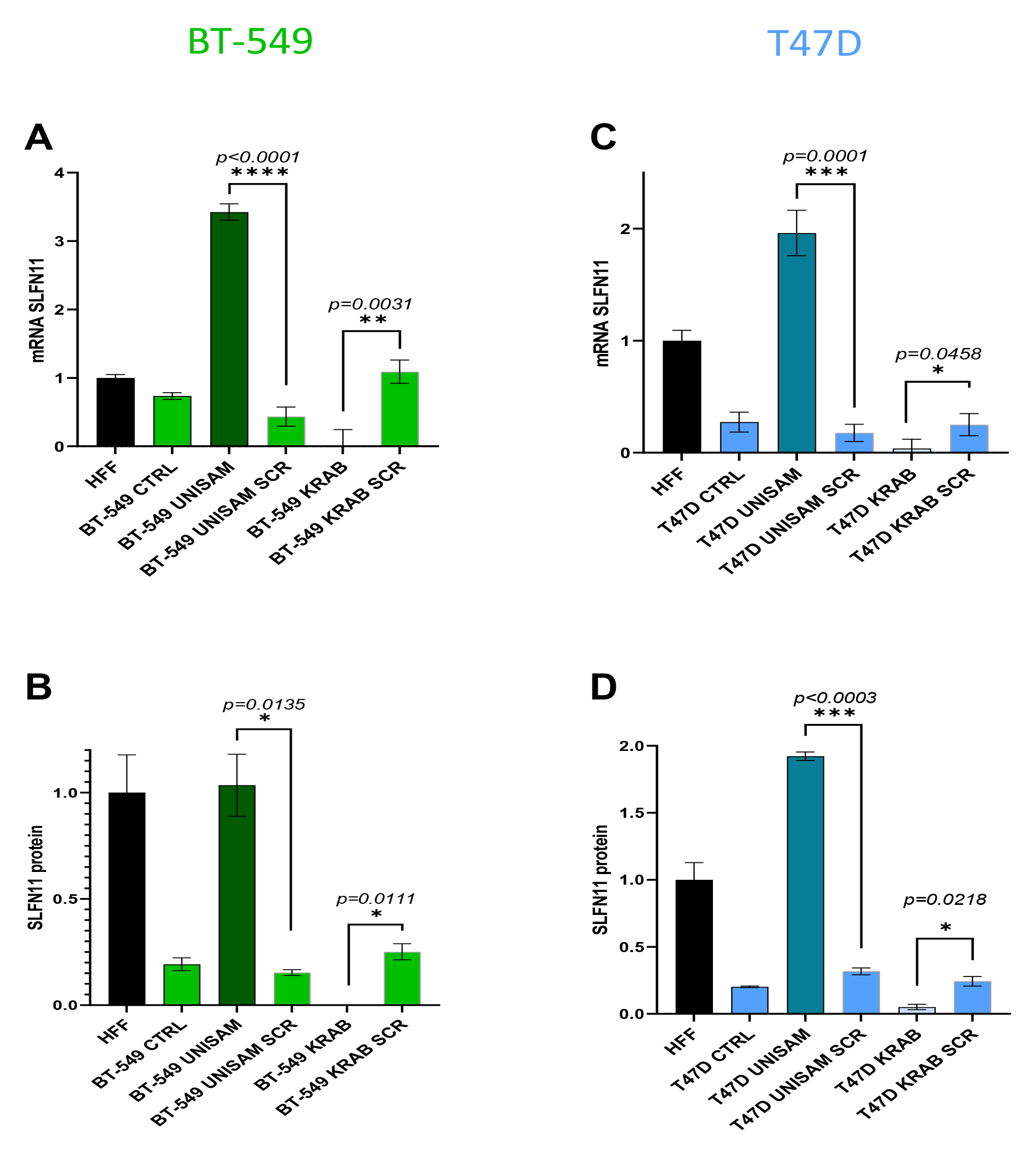
*CRISPR-dCas9 system significantly modulate SLFN11 expression* The UNISAM (unique Synergistic Activation Mediator) system developed by Fidanza, A., et al. was used for CRISPR activation of SLFN11 (Thakore et al. 2015) and KRAB (Krüppel associated box) was used for CRISPR inhibition of SLFN11 (Thakore et al. 2015). (A-B) Using the gRNA N7 we could successfully increase (with UNISAM) and decrease (with KRAB) SLFN11 expression in BT-549 as analyzed by Q-PCR (N=6) (A) and WES (N=4) (B). (C-D) Similarly, Using the gRNA N7 we could successfully increase (with UNISAM) and decrease (with KRAB) SLFN11 expression in T47D as analyzed by Q-PCR (N=6) (C) and WES (N=4) (D). In opposition, the UNISAM and KRAB system used with scramble gRNA (SCR) did not significantly affect SLFN11 expression compared to respective untreated cells (CTRL).

BT-549 and T47D cells were modified with the selected gRNA using UNISAM to increase SLFN11 expression and KRAB to decrease SLFN11 expression. Indeed, SLFN11 mRNA expression is significantly elevated in BT-549 UNISAM (p<0.0001) and significantly diminished in BT-549 KRAB (p =0.0031) when compared to respective scramble controls (SCR) (**Figure 3A**). Likewise, modulation of the SLFN11 protein level in BT-549 cells was significantly higher in UNISAM (p=0.0135) and lower in KRAB (p=0.0003) compared to respective scramble controls (**Figure 3B, Supplementary Figure 3H**). Similarly, T47D cells also showed a significant hike in SLFN11 mRNA expression using UNISAM (p=0.0001) and a decrease using KRAB (p=0.0458) in comparison to respective scramble controls (**Figure 3C**). Also, SLFN11 protein level was significantly increased in UNISAM (p=0.0003) and decreased in KRAB (p=0.0215) compared to their respective scrambled controls (**Figure 3D, Supplementary Figure 3I**). However, when working with MDA-MB- 231, despite an increase in SLFN11 mRNA expression **(Supplementary figure 2I**), no significant amount of protein could be detected in this strongly SLFN11 promoter methylated cell line using UNISAM **(Supplementary figure 3J**).

From these data, it is clear that CRISPR-UNISAM and CRISPR-KRAB combined with the appropriate gRNA protospacer (ACACTCGGACAGAATCCTGG or “N7”) can successfully increase and decrease endogenous SLFN11 expression in BT-549 and T47D cells. This system can modulate SLFN11 expression and is apparently the first report to demonstrate a stable system without inducing cell death.

### Modulation of SLFN11 expression sensitize the cells to Cisplatin, Epirubicin and Olaparib

The CRIPSR modified cells were then treated with different agents to assess the effect of SLFN11 expression on the sensitivity of cells to chemotherapeutic treatment. For each drugs treatment, different concentration ranges and timepoints were tested, depending on the drug at hand. For each dose response curves we assessed statistical significance at the concentration where nearly maximum effect was observed in the most sensitive cell line before toxicity plateaued (Figure 4 B, D, F, H, J, L).

**Figure 4:**
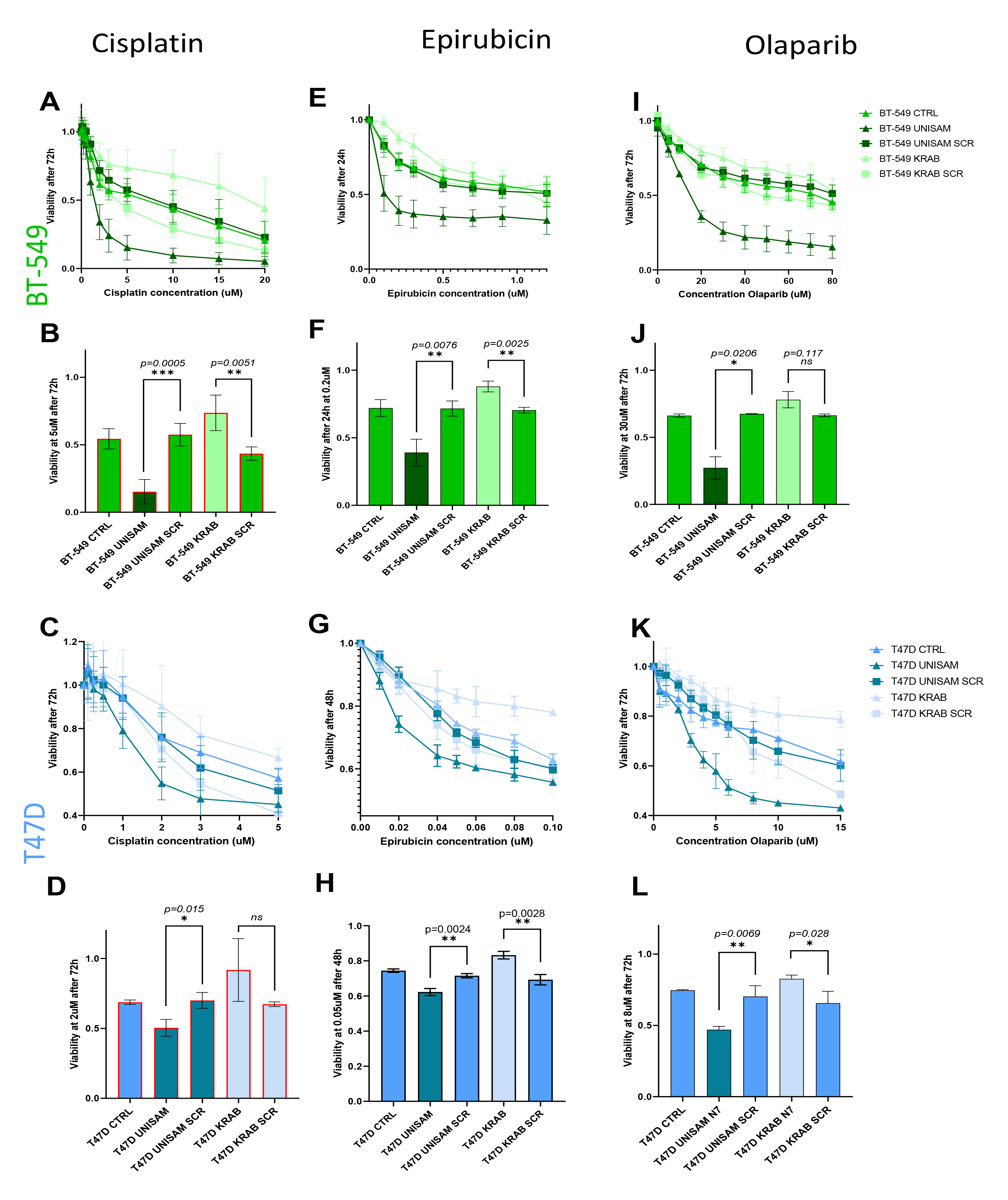
*CRISPR-dCas9 modulation of SLFN11 impact sensitivity to DNA Damaging agents* (A-B, E-F, H-I) In BT-549, SLFN11 increase with UNISAM (gRNA N7) or decrease with KRAB (gRNA N7) leads to respectively significant increase and decreased sensitivity to Cisplatin treatment (N=4) (A-B) Epirubicin treatment (E-F) (N=4) and Olaparib (N=4) (H- I) compared to respective scramble (SCR) controls. Only in Olapraib the decrease of SLFN11 with KRAB (gRNA N7) did not bring significant difference with SCR control at the indicated concentration. (C-D, G-H, J-K) In T47D, SLFN11 increase with UNISAM (gRNA N7) or decrease with KRAB (gRNA N7) leads to respectively significant increase and decreased sensitivity to Cisplatin treatment (N=4) (A-B) Epirubicin treatment (E-F) (N=3) and Olaparib (N=4) compared to respective scramble (SCR) controls. (J-K). Only in Cisplatin the decrease of SLFN11 with KRAB (gRNA N7) did not bring significant difference with SCR control at the indicated concentration.

Cisplatin, one of the most potent and widely used platinum-based drugs, was used to treat our modified BT-549 cells with a concentration ranging from 0.1 µM to 20 µM for 72h. After treatment cell viability was measured using ATP Lite and Calcein AM (**Figure 4A**). We observed that UNISAM modified cells, using the optimal gRNA (UNISAM), showed more sensitivity to cisplatin after 72h across a wide range of cisplatin concentrations compared to UNISAM using a scrambled gRNA (UNISAM SCR). In opposite, KRAB combined with this gRNA (KRAB) modified cells showed reduced sensitivity to cisplatin after 72h across a range of cisplatin concentration compared to KRAB used with scrambled gRNA (KRAB SCR). Indeed, at 5uM, BT-549 UNISAM exhibits significant increased sensitivity compared to UNISAM SCR (p=0.0005), while BT- 549 KRAB showed significant decrease sensitivity compared to KRAB SCR (p=0.0051) (**Figure 4B**).

Likewise, in modified T47D cells, upon treatment with Cisplatin for 72h we observed similar effect, with increased sensitivity of UNISAM modified cells and reduced sensitivity of KRAB modified cells across a range of concentration (**Figure 4C**). For example, at 2 µM for 72h, we can see a significant effect on drug sensitivity of T47D UNISAM cells compared to UNISAM SCR (p=0.015) (**Figure 4D**). On the other hand, T47D KRAB cells showed increased viability compared to scrambled although not significantly.

Epirubicin, belongs to the anthracycline family of chemotherapeutic drugs and is also commonly used for cancer treatment. We treated BT-549 cells with concentrations varying from 0.1 µM to 1.2 µM for 24h, and again observed increased sensitivity of UNISAM modified cells and reduced sensitivity of KRAB modified cells compared to respective controls (**Figure 4E**). For example, at 0.2 µM Epirubicin for 24h BT-549 UNISAM showed significant increase in sensitivity (p=0.0076) compared to UNISAM SCR, whereas BT-549 KRAB showed decrease sensitivity (p=0.0025) compared to KRAB SCR (**Figure 4F**).

T47D cells were also treated with Epirubicin with concentrations varying from 0.01 µM to0.1 µM for 48h, and again we observed increased sensitivity of UNISAM modified cells and reduced sensitivity of KRAB modified cells across a range of concentrations (**Figure 4G**). For example, at 0.05uM, T47D UNISAM showed significant increase in sensitivity (p=0.0024) compared to UNISAM SCR, whereas BT-549 KRAB showed decrease sensitivity compared to KRAB SCR (p=0.0028) (**Figure 4H**).

Olaparib is a Poly (adenosine diphosphate-ribose) polymerase inhibitor (PARPi) and is regarded as a promising anticancer agent. BT-549 cells were treated with Olaparib concentrations varying from 5 µM to 80 µM for 72h (**Figure 4I**). Once more, we observed increased sensitivity of UNISAM modified cells and reduced sensitivity of KRAB modified cells in a range of concentration. For instance, at 30 µM, BT-549 UNISAM shows increased sensitivity (p=0.0206) and BT-549 KRAB showed decreased sensitivity but not significantly compared to relevant scramble controls (**Figure 4J**). Similarly, T47D cells were treated with Olaparib at concentrations ranging from 0.5 µM to 15 µM for 72h (**Figure 4K**). Once more, UNISAM modified cells displayed increased sensitivity KRAB modified cells showed reduced sensitivity compared to their controls. Like at 8 µM, and compared to respective scrambled controls, T47D UNISAM displayed increased sensitivity (p=0.0069), while T47D KRAB showed decreased sensitivity (p=0.028) (**Figure 4L**).

In conclusion, CRISPR-UNISAM and CRISPR-KRAB systems used with appropriate gRNA efficiently increase and decrease SLFN11 expression in BT-549 and T47D which in turn modulates the sensitivity to Cisplatin, Epirubicin, and Olaparib. We can infer that modulation of SLFN11 expression can effectively impact the effect of DNA damaging agents on these cell lines.

### RNAseq analysis of CRISPR modified cells

To further comprehend the mechanism of increased resistance and sensitivity of the CRISPR modified cell lines to cisplatin treatment, we performed RNA sequencing of our modified cells prior and after Cisplatin treatment. Our RNA sequencing was done in triplicate on three independent batches of cells resulting in 5-9 replicates for each condition. The batch effects observed resulting from cell culture and RNA isolation was identified by PCA analysis (**Supplementary Figure 4**). This batch effect was then resolved using Combat (Leek et al. 2012).

PCA analysis clearly shows a separation between the Cisplatin treated and non-treated cells. Indeed, while the CRISPR modified cells clustered close to unmodified or scramble modified cells, the Cisplatin treatment of cells resulted in a drastic shift of the treated cells (represented by the red arrows in figure 5A). Under Cisplatin treatment the effect of the CRISPR modification between UNISAM (shift indicated in pink), KRAB (shift indicated in orange) and unmodified cells seem to be much larger and captured by a different dimension in the PCA plot.

**Figure 5:**
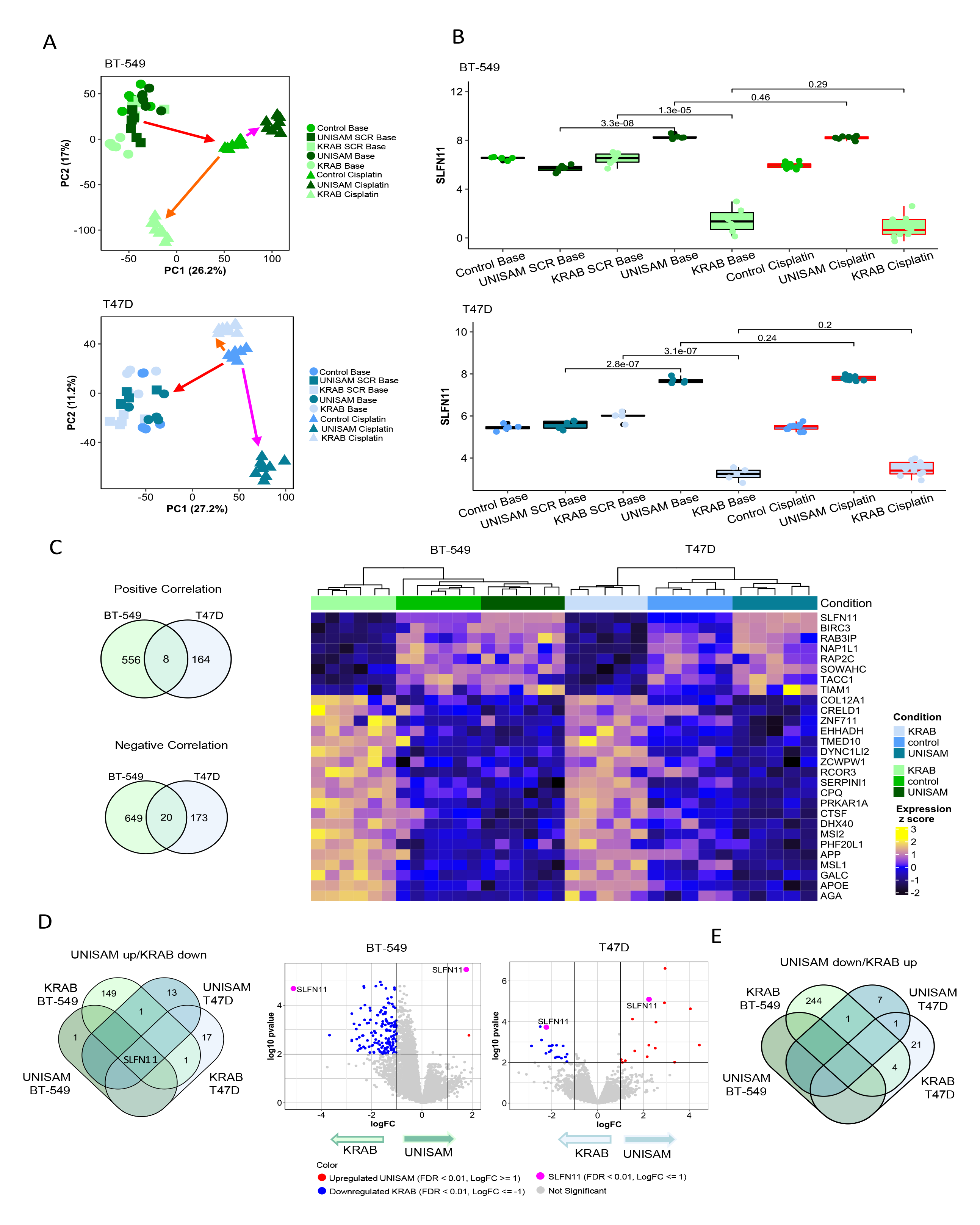
*RNAseq analysis of CRISPR modified cells* (A). Principal component analysis based on gene expression for all samples (baseline and cisplatin-treated samples) post performing combat on samples to remove batch effect for BT-549 and T47D cell lines. (B). Expression of SLFN11 across baseline and cisplatin-treated samples in BT-549 and T47D cell lines. (C). VennDiagram of common genes between BT-549 and T47D which are positively (N = 8) and negatively (N = 20) correlating with SLFN11 (correlation coefficient R < |0.5|) based on the baseline samples. Heatmap showing the expression of positively and negatively correlating genes with SLFN11 (N = 28). (D). VennDiagram of differentially expressed genes between BT-549 and T47D cell lines in UNISAM up (vs. control) and KRAB down (vs. control) (FDR < 0.01, logFC < |1|). Volcano plot showing upregulated DEG between UNISAM vs, control and downregulated DEG between KRAB vs, control in baseline samples (FDR < 0.01, logFC < |1| considered as significant). (E). VennDiagram of differentially expressed genes between BT-549 and T47D cell lines in UNISAM down (vs. control) and KRAB up (vs. control) (FDR < 0.01, logFC < |1|) in baseline samples.

In alignment with our Q-PCR and western blot analysis, RNA sequencing confirmed that UNISAM modification of both BT-549 and T47D resulted in a strong increase of SLFN11 RNA expression in baseline (p=3,3.10^-8^ and p=2,8.10^-7^ respectively), while KRAB significantly reduced the expression of SLFN11 in baseline (p=1,3.10^-5^ and p=3,1.10^-7^ respectively). Of note, Cisplatin treatment did not affect SLFN11 expression in UNISAM in BT-549 and T47D (p = 0.46 and p = 0.24 respectively) as well in KRAB (p = 0.29 and p = 0.2 respectively) when compared to corresponding baseline samples (**Figure 5B**).

To appreciate the consistency of the perturbations caused by SLFN11 modulation across cell lines we performed Pearson correlation and analyzed the genes that were positively or negatively correlated (R>+/- 0.5) with SLFN11 expression in both UNISAM and KARB modified BT-549 and T47D cell lines. Among those genes 8 were positively and 20 were negatively correlated with SLFN11 in both cell lines (**figure 5C**).

The analysis of differentially expressed genes (DEG) using a log fold change ≥1 and FDR ≥ 0.01, showed that only a few genes were upregulated or down regulated along with SLFN11 in UNISAM and KRAB in each cell lines compared to controls modified with a scrambled guide RNA **(volcano plots in figure 5D**). Though, only SLFN11 was commonly up regulated in UNISAM and downregulated in KRAB between the two cell lines (**Figure 5D**). No common genes between the two cell lines could be identified as down regulated in UNISAM and upregulated in KRAB (**Figure 5E**). These data confirm the specificity of the UNISAM and KRAB CRISPR systems for up or down regulating SLFN11 specifically.

### RNASeq analysis of CRISPR modified cells treated with Cisplatin

To identify genes associated with SLFN11 up and down-regulation under Cisplatin treatment in both cell lines, we performed DEG analysis comparing UNISAM and KRAB modified cells with unmodified controls, using Limma and considered genes with log fold change ≥1 and FDR ≥ 0.01 as significant. A list of DEGs comparing cells modified with scrambled gRNA’s vs unmodified cells was used to remove false positives. When only considering genes affected in the same way in both cell lines there are 92 genes upregulated in UNISAM / downregulated in KRAB under cisplatin treatment (red). On the other hand, 80 genes were found to be downregulated in UNISAM / upregulated in KRAB under cisplatin treatment (green) (**Figure 6A**).

**Figure 6:**
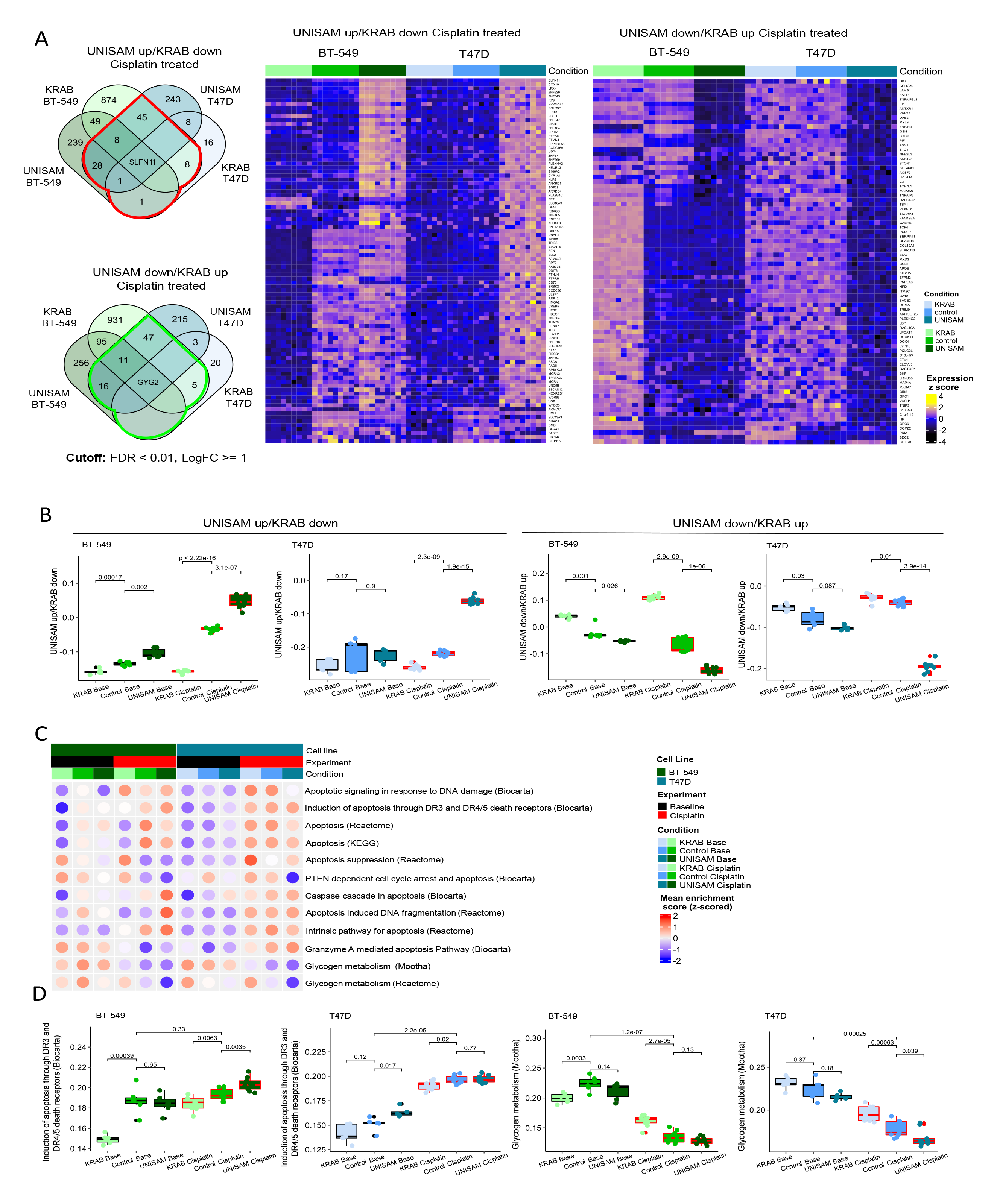
*RNASeq analysis of CRISPR modified cells treated with Cisplatin* (A). VennDiagram of differentially expressed genes between BT-549 and T47D cell lines in UNISAM up/KRAB down (vs. control), and UNISAM down/KRAB up (vs. control) respectively, (FDR < 0.01, logFC < |1|) in Cisplatin-treated samples. Heatmaps illustrate the expression of DEG genes in UNISAM up/KRAB down and UNISAM down/KRAB up (genes highlighted in red and green in VennDiagram, respectively). (B). Boxplot of enrichment score generated for list of DEG (UNISAM up/KRAB down (n = 94) and UNISAM down/KRAB up (n = 87)) in BT-549 and T47D cell lines. (C). Dotted heatmap of mean enrichment score for apoptosis and glycogen metabolism pathways. (D). Boxplot of enrichment scores for selected apoptosis and glycogen metabolism pathways in BT- 549 and T47D cell lines.

Enrichment score (ES) of genes upregulated in UNISAM / downregulated in KRAB (n=92) and genes downregulated in UNISAM / upregulated in KRAB (n=80) was calculated using ssGSEA to create a SLFN11 signature for genes that go in the same or opposite direction as SLFN11, respectively. As expected under cisplatin treatment this upregulated in UNISAM / downregulated in KRAB score is low in KRAB, high in UNISAM with the control cells in the middle and the reverse seen in the when looking at the downregulated in UNISAM / upregulated in KRAB score. What is interesting is that the same pattern, although much weaker is observed in the cell not treated with cisplatin, indicating the change in DNA Damage response machinery. (**Figure 6B**).

Next, we aimed to identify the enriched pathways using genes from the DEG analysis. The above-mentioned gene lists were uploaded to Ingenuity Pathway Analysis (IPA) database. Results show that there are 22 pathways that show enrichment of genes in these gene lists (**Figure S5**). However, we could not identify pathways associated with cell death and apoptosis as one would expect. This incentivized us to perform a more guided pathway analysis by looking specifically at apoptosis pathways, in addition to glycogen metabolic pathways since we found that GYG2 was consistently downregulated in UNISAM and upregulated in KRAB in both BT-549 and T47D cell lines treated with Cisplatin. To do so, we downloaded gene sets belong to these pathways from MSigDB and calculated an ES for each pathway and visualized these in a heatmap (**Figure 6C**). As expected, enrichment scores show that most apoptosis pathways have higher enrichment in Cisplatin treated samples compared to baseline, and the reverse is observed for glycogen metabolism pathways. To Illustrate the effect SLFN11 has in this context, we selected the pathways showing the clearest trend in both cell lines (**Figure 6D**). Obviously, higher enrichment of apoptosis pathway is observed in Cisplatin treated samples compared to control. More interestingly, within Cisplatin treated sample, apoptosis is significantly more enriched in UNISAM and less enriched in KRAB in BT-549 (p = 0.0035, p = 0.0063 respectively), however in T47D enrichment was only significant higher in KRAB (p = 0.02) when compared to corresponding controls. Whereas for glycogen metabolism pathway, lower enrichment was observed in Cisplatin treated samples compared to baseline samples. Glycogen metabolism is significantly higher enriched in KRAB in BT-549 and T47D (p = 2,7.10^-5^, p = 0.00063 respectively), whereas it was significantly less enriched in UNISAM in T47D only (p = 0.039) compared to control. This would be in line with the higher energetic requirements of a proliferating cell or a cell with active DNA repair machinery. (Sobanski et al. 2021)

## Discussion

Since the 2012 discovery of the role of SLFN11 as prognostic marker for response to DDAs in-silico (Zoppoli et al. 2012). It was demonstrated that the reduction of SLFN11 expression led to increased resistance to such treatment in-vitro, due to its reduced ability to inhibit replication checkpoint maintenance and HR repair upon DNA damage (Mu et al. 2016). This is further substantiated by the predictive value of SLFN11 immunohistochemistry in ovarian cancer treated with Cisplatin.(Winkler et al. 2021) It was therefore postulated that the opposite should also be true. Unfortunately, in our hands traditional over expression of SLFN11 in cell lines resulted in cell death (data not shown). We therefore investigated alternative ways to increase SLFN11 expression. It was previously established that SLFN11 was an IFN inducible gene (Mavrommatis, Fish, and Platanias 2013), and that methylation was the primary mechanism of regulation of SLFN11 (Nogales et al. 2015; Peng et al. 2019). Indeed, in our *in vitro* setting we could establish a good correlation between SLFN11 promoter methylation and gene expression or protein expression. Also, the use of IFN-γ and a well-established demethylation agent (DAC) could indeed increase SLFN11 expression *in vitro*. Though, this increase despite being statistically significant, was limited. We therefore attempted specific increase in SLFN11 expression using a dCas9 CRISPR activation system. We clearly observed in repeated experiments with multiple drugs and in multiple cell lines that this CRISPR system leads to a strong upregulation of SLFN11 leading in turn to increased sensitivity of those cell lines to DDA treatment or Topo isomerase inhibitor treatment. On the flip side, the down regulation using a dCas9 CRISPR inhibition system led to increased resistance to similar treatments in the same cell lines. By transcriptomic analysis we could establish the specificity of those CRISPR systems used to up and down regulate SLFN11 as the only gene upregulated by the UNISAM system in both T47D and BT-549 was SLFN11, and in the reverse context, the only gene identified as down-regulated by KRAB system in both T47D and BT-549 was SLFN11. Therefore, we can be confident that the observed effects on drugs sensitivity where specifically due to SLFN11 modulation.

We should note that we observed these changes in chemo sensitivity in cell lines which still exhibit basal expression of SLFN11 and only moderate methylation of its promoter (50% methylation in the case of T47D), but in some cell lines such as MDA-MB-231, with strong repression of SLFN11 expression *via* complete promoter methylation our dCas9 CRISPR activation system could not increase significantly SLFN11 expression. The DAC treatment, on the other hand, resulted in a partial demethylation of the promoter. We believe that a cumulation of different strategies could therefore be key. Indeed, while increasing SLFN11 expression can be beneficial in many clinical cases, increasing the expression in patients with no SLFN11 expression might be more essential. Therefore, it is worth testing *in-vivo* if the combined treatment with previously approved drugs like demethylation agents such as DAC combined with immune checkpoint blockade would lead to sufficient increase in SLFN11 expression in tumors with a decreased SLFN11 expression. If using these existing agents would not meaningfully improve chemosensitivity, more potent approaches like an *in-vivo* application of the dCAS9 method we used in-vitro could be evaluated. We believe in dept exploration of the feedback loop observed between tumor cells SLFN11 expression and T-cells IFN-γ expression is required to understand the DNA damage response mechanisms interactions with the anti-tumor immune response. (Mezzadra et al. 2019; Winkler et al. 2021). Also, while clinical phase I trials using CRISPR are on the rise, the use of CRISPR systems such as activation system require permanent expression in target cell and could lead to additional advers effect due to integration of the constructs. Alternatively, other dCAS CRISPR systems allow to specifically demethylate genes promoters. As methylation is key to unlocking SLFN11 expression, it would be worth investigating if in such resistant cell lines, CRISPR demethylation would be sufficient to stably increase SLFN11 expression alone or in combination with other treatments.

---

*Supplementary figure 1: gRNA location and CRISPR modification*

(A) The predicted promoter region of SLFN11 (in green) is surrounding the gene’s exon1 and CpG island (in red) analysis show its location in the center of the promoter area. gRNAs were therefore designed along the central region of the promoter of SLFN11 (N1 to N10). (B) schematic representation of the strategy adopted to respectively increase SLFN11 expression using UNISAM system and decrease SLFN11 expression using KRAB system. After insertion of the gRNA into the respective plasmids, cells were transformed with the integrative plasmids using electroporation and selected for the expression of respectively mCherry or GFP reporter genes. Cells were then analyzed for SLFN11 expression by westernblot and by Q-PCR.

*Supplementary figure 2: Screening of gRNA efficiency at upregulating or downregulating SLFN11 using UNISAM or KRAB systems*

(A-D) Relative mRNA expression of SLFN11 analyzed by RT-qPCR (n=3) (A-C) and relative SLFN11 protein expression analyzed by WES (n=2) (B-D) in BT-549 cancer cell lines modified with each gRNA for CRISPR-dCas9-UNISAM (A-B) or CRISPR-dCas9-KRAB (C-D) relative to non-modified cell line.

(E-H) Relative mRNA expression of SLFN11 analyzed by RT-qPCR (n=3) (E-G) and relative SLFN11 protein expression analyzed by WES (n=2) (F-H) in T47D cancer cell lines modified with each gRNA for CRISPR-dCas9-UNISAM (E-F) or 5 (N1, N2, N6, N7 and N10) of the 7 gRNA for CRISPR-dCas9-KRAB (G-H) relative to non-modified cell line.

(I) Relative mRNA expression of SLFN11 analyzed by RT-qPCR in MDA-MB-231 cancer cell lines modified with each gRNA for CRISPR-dCas9-UNISAM relative to non-modified cell line.

*Supplementary figure 3: representative WES results*

(A) Representative WES results of the analysis of SLFN11 protein expression in the 8 tested unmodified breast cancer cell lines compared to HFF. (B) Representative SLFN11 protein expression in BT-549, T47D and MDA-MB-231 analyzed by WES after treatment with 5uM of DAC for 72h compared to the expression level in untreated HFF. (C) Representative SLFN11 protein expression in BT-549, T47D and MDA-MB-231 analyzed by WES after treatment with 5nM of IFN-γ for 24h compared to the expression level in untreated HFF. (D-E) Representative SLFN11 protein expression in BT-549 (D) or T47D (E) modified with UNISAM and each of the 7 gRNA compared to the respective unmodified cells. (F-G) Representative SLFN11 protein expression in BT-549 (F) or T47D (G) modified with KRAB and each of the 7 gRNA compared to the respective unmodified cells. (H-I) Representative SLFN11 protein expression level analyzed by WES in BT-549 (H) or T47D (I) after modification with UNISAM (gRNA 7 or gRNA SCR) or KRAB (gRNA 7 or gRNA SCR) compared to respective unmodified cells and HFF. (J) Representative SLFN11 protein expression level analyzed by WES in MDA-MB-231 after modification with UNISAM (gRNA 7) compared to respective unmodified cells and HFF.

*Supplementary figure 4: Principal component analysis pre and post-combat*

Principal component analysis based on gene expression for all samples (baseline and cisplatin-treated samples) pre and post performing combat on samples to remove batch effect for BT-549 and T47D cell lines.

*Supplementary figure 5: Pathway enrichment analysis*

Enriched pathways associated with DEG (n = 181, FDR < 0.01, LogFC >= 1) from limma analysis in UNISAM up/KRAB down and UNISAM down/KRAB up, using Ingenuity Pathway Analysis (IPA).

## Notes

### Competing Interest Statement

The authors have declared no competing interest.

## References

Anders, Simon, Paul Theodor Pyl, and Wolfgang Huber. 2015. “HTSeq—a Python Framework to Work with High-Throughput Sequencing Data.” Bioinformatics 31(2): 166–69. https://doi.org/10.1093/bioinformatics/btu638.

Barretina, J., G. Caponigro, N. Stransky, K. Venkatesan, A.A. Margolin, S. Kim, C.J. Wilson, et al. 2012. “22 The Cancer Cell Line Encyclopedia - Using Preclinical Models to Predict Anticancer Drug Sensitivity.” European Journal of Cancer 48 (July): S5–6. https://doi.org/10.1016/S0959-8049(12)70726-8.

Bolstad, Ben. 2016. “PreprocessCore: A Collection of Pre-Processing Functions Version 1.36 from Bioconductor.” 2016. https://github.com/bmbolstad/preprocessCore.

Chen, Hanbo, and Paul C. Boutros. 2011. “VennDiagram: A Package for the Generation of Highly-Customizable Venn and Euler Diagrams in R.” BMC Bioinformatics 12(1): 35. https://doi.org/10.1186/1471-2105-12-35.

Dobin, Alexander, Carrie A. Davis, Felix Schlesinger, Jorg Drenkow, Chris Zaleski, Sonali Jha, Philippe Batut, Mark Chaisson, and Thomas R. Gingeras. 2013. “STAR: Ultrafast Universal RNA-Seq Aligner.” *Bioinformatics (Oxford*, England) 29 (1): 15–21. https://doi.org/10.1093/bioinformatics/bts635.

Fidanza, Antonella, Martha Lopez-Yrigoyen, Nicola Romanò, Rhiannon Jones, A. Helen Taylor, and Lesley M. Forrester. 2017. “An All-in-One UniSam Vector System for Efficient Gene Activation.” Scientific Reports 7 (1): 6394. https://doi.org/10.1038/s41598-017-06468-6.

Gu, Zuguang, Lei Gu, Roland Eils, Matthias Schlesner, and Benedikt Brors. 2014. “Circlize Implements and Enhances Circular Visualization in R.” Bioinformatics 30(19): 2811–12. https://doi.org/10.1093/bioinformatics/btu393.

Hänzelmann, Sonja, Robert Castelo, and Justin Guinney. 2013. “GSVA: Gene Set Variation Analysis for Microarray and RNA-Seq Data.” BMC Bioinformatics 14 (January): 7. https://doi.org/10.1186/1471-2105-14-7.

Isnaldi, Edoardo, Domenico Ferraioli, Lorenzo Ferrando, Sylvain Brohée, Fabio Ferrando, Piero Fregatti, Davide Bedognetti, Alberto Ballestrero, and Gabriele Zoppoli. 2019. “Schlafen-11 Expression Is Associated with Immune Signatures and Basal-like Phenotype in Breast Cancer.” Breast Cancer Research and Treatment 177 (2): 335–43. https://doi.org/10.1007/s10549-019-05313-w.

Leek, Jeffrey T., W. Evan Johnson, Hilary S. Parker, Andrew E. Jaffe, and John D. Storey. 2012. “The Sva Package for Removing Batch Effects and Other Unwanted Variation in High-Throughput Experiments.” Bioinformatics 28 (6): 882–83. https://doi.org/10.1093/bioinformatics/bts034.

Li, Manqing, Elaine Kao, Xia Gao, Hilary Sandig, Kirsten Limmer, Mariana Pavon-Eternod, Thomas E. Jones, et al. 2012. “Codon-Usage-Based Inhibition of HIV Protein Synthesis by Human Schlafen 11.” Nature 491 (7422): 125–28. https://doi.org/10.1038/nature11433.

Liberzon, A., A. Subramanian, R. Pinchback, H. Thorvaldsdottir, P. Tamayo, and J. P. Mesirov. 2011. “Molecular Signatures Database (MSigDB) 3.0.” Bioinformatics 27(12): 1739–40. https://doi.org/10.1093/bioinformatics/btr260.

Mavrommatis, Evangelos, Eleanor N. Fish, and Leonidas C. Platanias. 2013. “The Schlafen Family of Proteins and Their Regulation by Interferons.” Journal of

Interferon & Cytokine Research 33 (4): 206–10. https://doi.org/10.1089/jir.2012.0133.

Mezzadra, Riccardo, Marjolein de Bruijn, Lucas T. Jae, Raquel Gomez-Eerland, Anja Duursma, Ferenc A. Scheeren, Thijn R. Brummelkamp, and Ton N. Schumacher. 2019. “SLFN11 Can Sensitize Tumor Cells towards IFN-γ-Mediated T Cell Killing.” PLoS ONE 14 (2). https://doi.org/10.1371/journal.pone.0212053.

Mu, Yanhua, Jiangman Lou, Mrinal Srivastava, Bin Zhao, Xin-hua Feng, Ting Liu, Junjie Chen, and Jun Huang. 2016. “SLFN11 Inhibits Checkpoint Maintenance and Homologous Recombination Repair.” EMBO Reports 17 (1): 94–109. https://doi.org/10.15252/embr.201540964.

Murai, Junko, Sai-Wen Tang, Elisabetta Leo, Simone A. Baechler, Christophe E. Redon, Hongliang Zhang, Muthana Al Abo, et al. 2018. “SLFN11 Blocks Stressed Replication Forks Independently of ATR.” Molecular Cell 69 (3): 371–384.e6. https://doi.org/10.1016/j.molcel.2018.01.012.

Murai, Junko, Hongliang Zhang, Lorinc Pongor, Sai-Wen Tang, Ukhyun Jo, Fumiya Moribe, Yixiao Ma, Masaru Tomita, and Yves Pommier. 2020. “Chromatin Remodeling and Immediate Early Gene Activation by SLFN11 in Response to Replication Stress.” Cell Reports 30 (12): 4137–4151.e6. https://doi.org/10.1016/j.celrep.2020.02.117.

Nogales, Vanesa, William C. Reinhold, Sudhir Varma, Anna Martinez-Cardus, Catia Moutinho, Sebastian Moran, Holger Heyn, et al. 2015. “Epigenetic Inactivation of the Putative DNA/RNA Helicase SLFN11 in Human Cancer Confers Resistance to Platinum Drugs.” Oncotarget 7 (3): 3084–97. https://doi.org/10.18632/oncotarget.6413.

Parsi, Krishna Mohan, Erica Hennessy, Nicola Kearns, and René Maehr. 2017. “Using an Inducible CRISPR-DCas9-KRAB Effector System to Dissect Transcriptional Regulation in Human Embryonic Stem Cells.” In Eukaryotic Transcriptional and Post-Transcriptional Gene Expression Regulation, edited by Narendra Wajapeyee and Romi Gupta, 1507:221–33. Methods in Molecular Biology. New York, NY: Springer New York. https://doi.org/10.1007/978-1-4939-6518-2_16.

Peng, Yaojun, Li Wang, Liangliang Wu, Ling Zhang, Guangjun Nie, and Mingzhou Guo. 2019. “Methylation of SLFN11 Promotes Gastric Cancer Growth and Increases Gastric Cancer Cell Resistance to Cisplatin.” Journal of Cancer 10 (24): 6124–34. https://doi.org/10.7150/jca.32511.

Risso, Davide, Katja Schwartz, Gavin Sherlock, and Sandrine Dudoit. 2011. “GC-Content Normalization for RNA-Seq Data.” BMC Bioinformatics 12 (1): 480. https://doi.org/10.1186/1471-2105-12-480.

Ritchie, Matthew E., Belinda Phipson, Di Wu, Yifang Hu, Charity W. Law, Wei Shi, and Gordon K. Smyth. 2015. “Limma Powers Differential Expression Analyses for RNA-Sequencing and Microarray Studies.” Nucleic Acids Research 43 (7): e47– e47. https://doi.org/10.1093/nar/gkv007.

Sobanski, Thais, Maddison Rose, Amila Suraweera, Kenneth O’Byrne, Derek J. Richard, and Emma Bolderson. 2021. “Cell Metabolism and DNA Repair Pathways: Implications for Cancer Therapy.” Frontiers in Cell and Developmental Biology 9 (March): 633305. https://doi.org/10.3389/fcell.2021.633305.

Stewart, C. Allison, Pan Tong, Robert J. Cardnell, Triparna Sen, Lerong Li, Carl M. Gay, Fatemah Masrorpour, et al. 2017. “Dynamic Variations in Epithelial-to-Mesenchymal Transition (EMT), ATM, and SLFN11 Govern Response to PARP Inhibitors and Cisplatin in Small Cell Lung Cancer.” Oncotarget 8 (17): 28575–87. https://doi.org/10.18632/oncotarget.15338.

Tang, Sai-Wen, Anish Thomas, Junko Murai, Jane B. Trepel, Susan E. Bates, Vinodh N. Rajapakse, and Yves Pommier. 2018. “Overcoming Resistance to DNA-Targeted Agents by Epigenetic Activation of Schlafen 11 ( *SLFN11)* Expression with Class I Histone Deacetylase Inhibitors.” Clinical Cancer Research 24 (8): 1944–53. https://doi.org/10.1158/1078-0432.CCR-17-0443.

Thakore, Pratiksha I., Anthony M. D’Ippolito, Lingyun Song, Alexias Safi, Nishkala K. Shivakumar, Ami M. Kabadi, Timothy E. Reddy, Gregory E. Crawford, and Charles A. Gersbach. 2015. “Highly Specific Epigenome Editing by CRISPR-Cas9 Repressors for Silencing of Distal Regulatory Elements.” Nature Methods 12 (12): 1143–49. https://doi.org/10.1038/nmeth.3630.

Wickham, Hadley. 2009. Ggplot2: Elegant Graphics for Data Analysis. Use R! New York: Springer-Verlag. https://doi.org/10.1007/978-0-387-98141-3.

Winkler, Claudia, Matthew King, Julie Berthe, Domenico Ferraioli, Anna Garuti, Federica Grillo, Jaime Rodriguez-Canales, et al. 2021. “SLFN11 Captures Cancer-Immunity Interactions Associated with Platinum Sensitivity in High-Grade Serous Ovarian Cancer.” JCI Insight 6 (18): e146098. https://doi.org/10.1172/jci.insight.146098.

Zoppoli, Gabriele, Marie Regairaz, Elisabetta Leo, William C. Reinhold, Sudhir Varma, Alberto Ballestrero, James H. Doroshow, and Yves Pommier. 2012. “Putative DNA/RNA Helicase Schlafen-11 (SLFN11) Sensitizes Cancer Cells to DNA-Damaging Agents.” Proceedings of the National Academy of Sciences of the United States of America 109 (37): 15030–35. https://doi.org/10.1073/pnas.1205943109.

