## Supplementary figures for "Modulation of SLFN11 induces changes in DNA Damage response"

### Supplementary figure 1

**A**

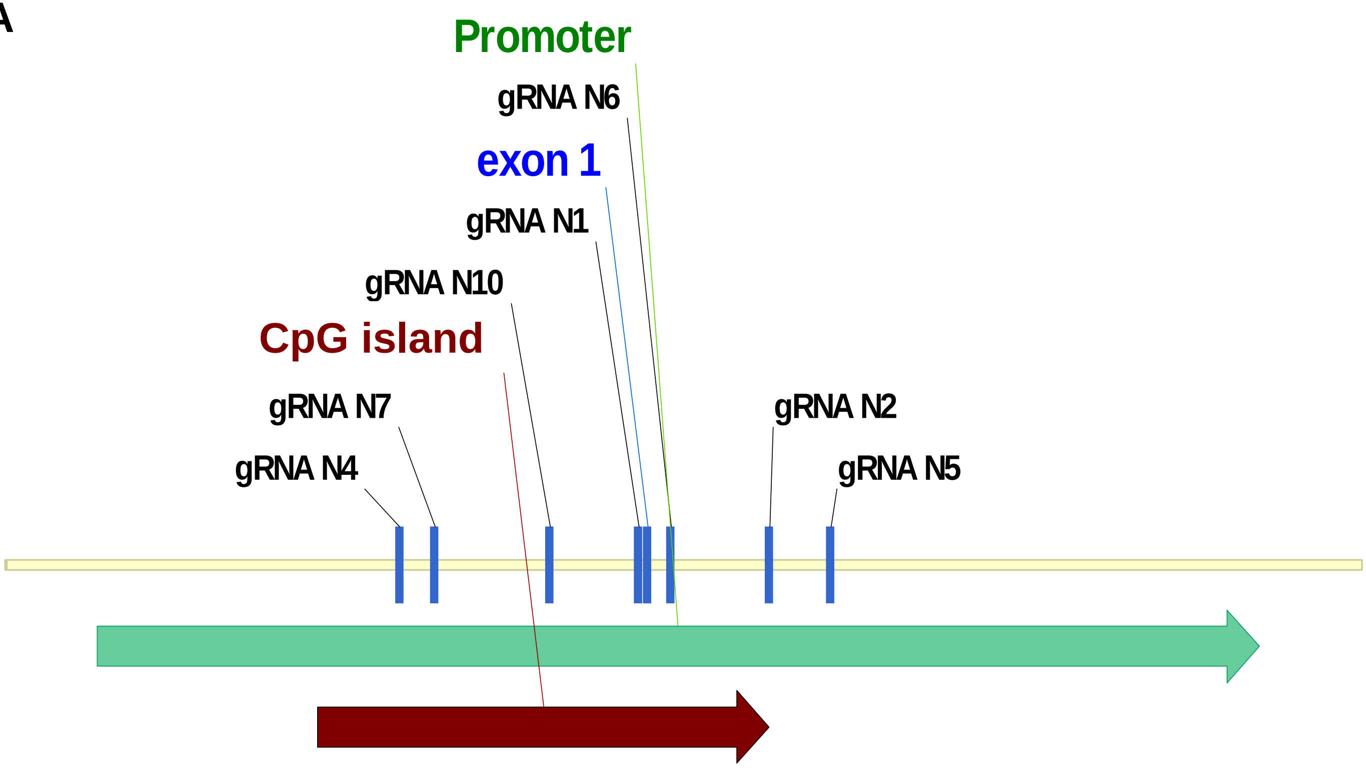


**B**

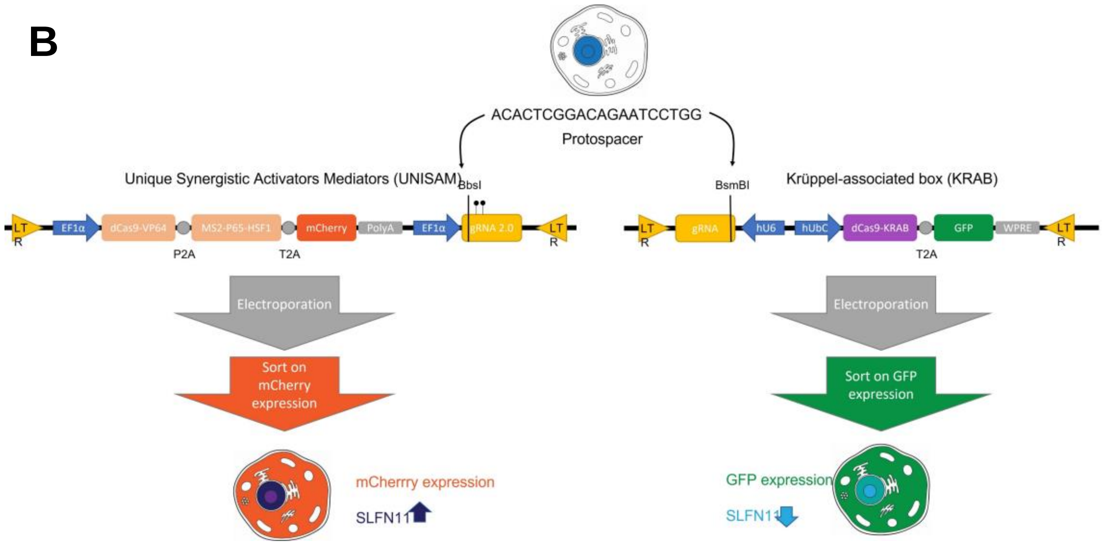

### Supplementary figure 2

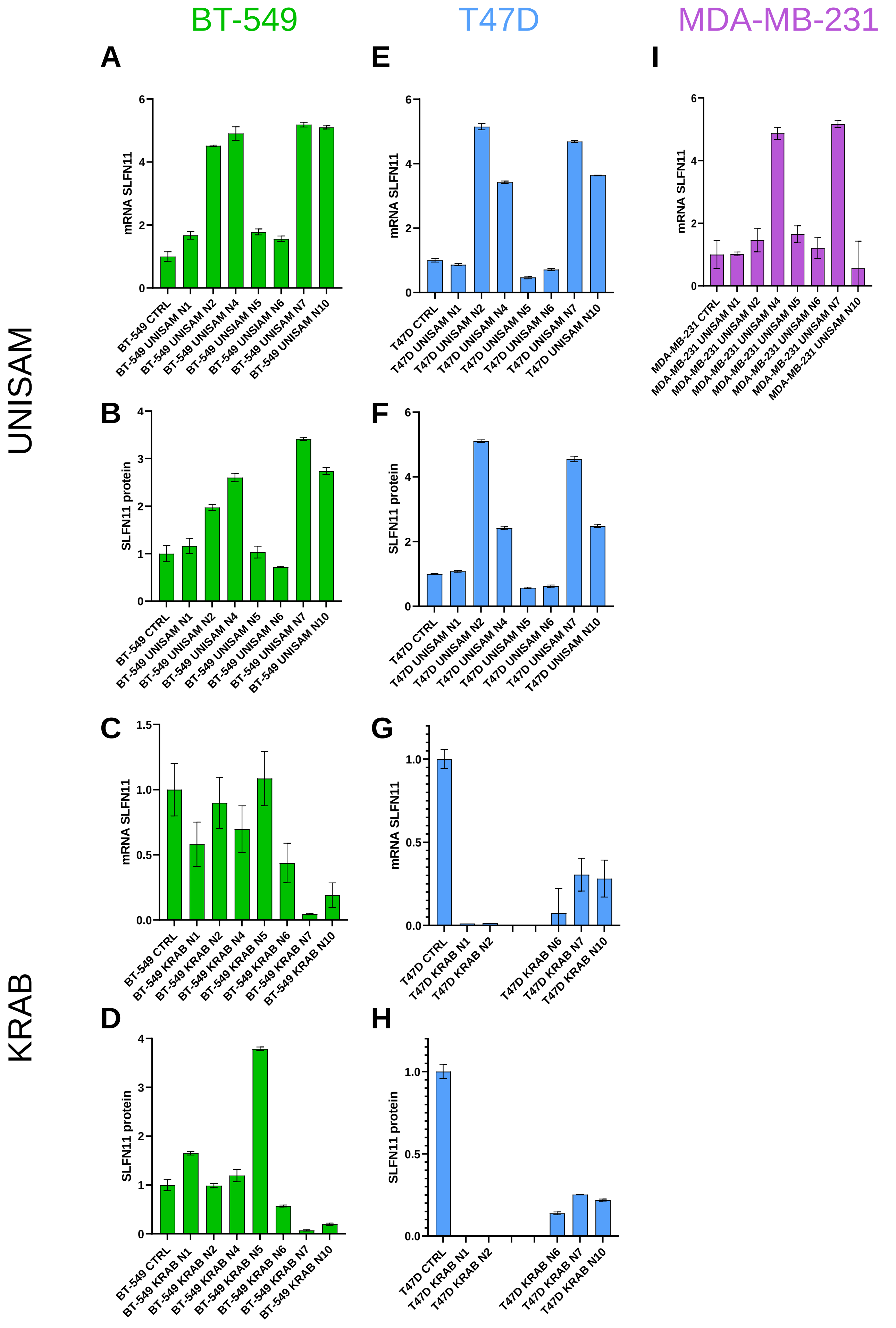

### Supplementary figure 3

**A** **Baseline**

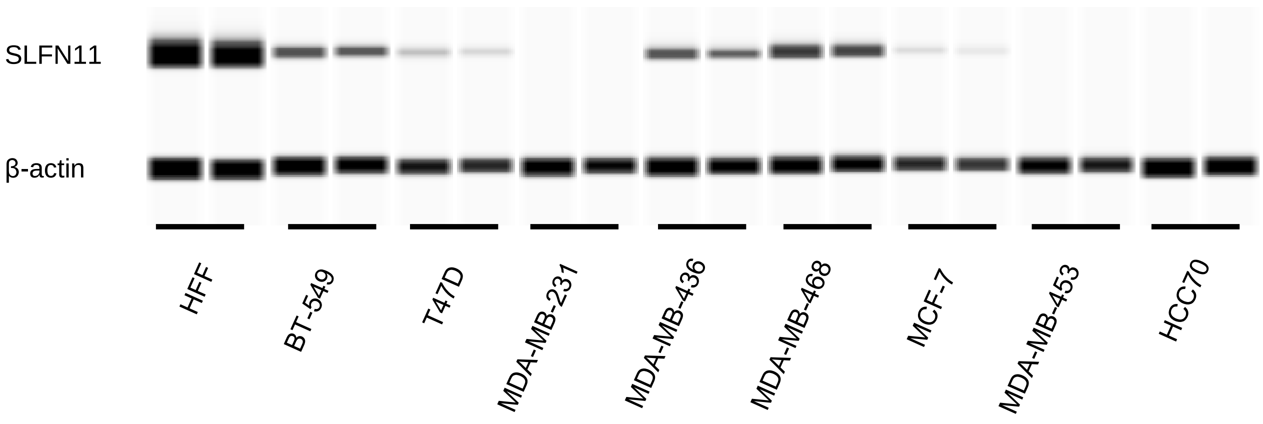


**B** **Decitabine treatment**

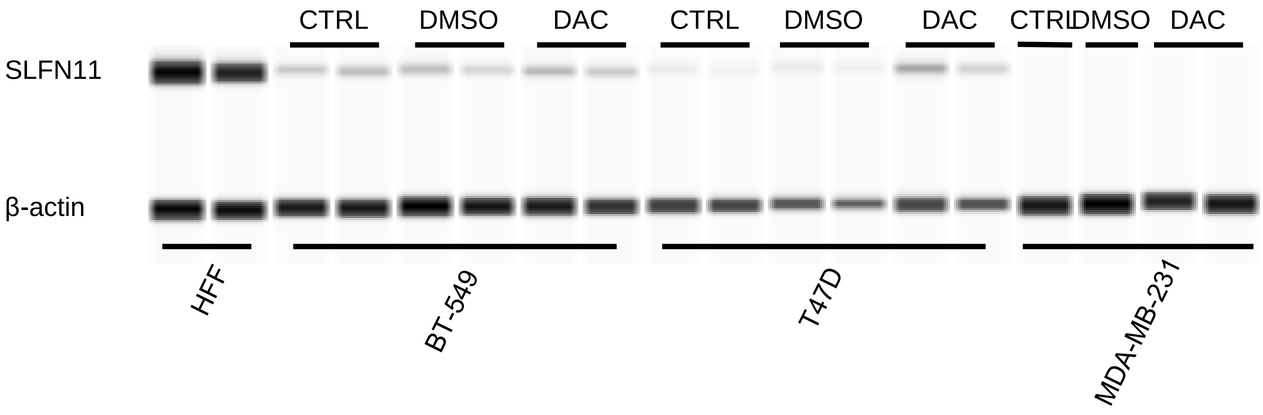


**C** **IFN-γ treatment**

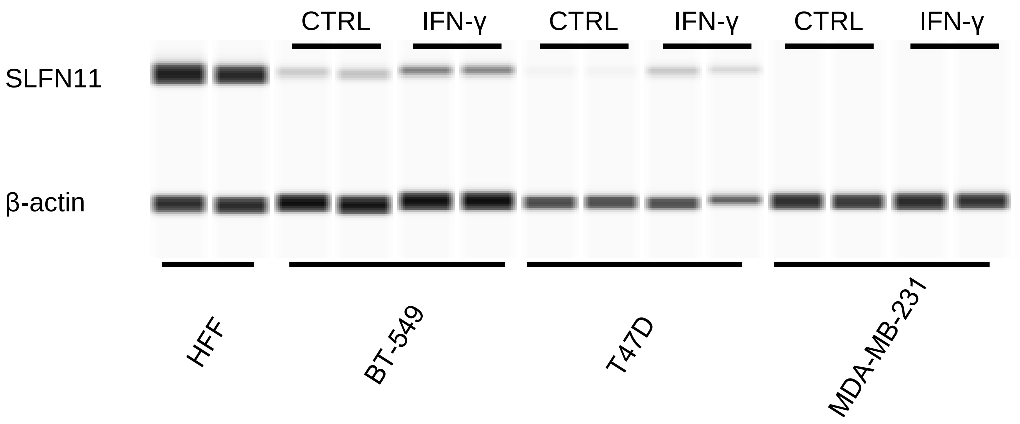

**D** **BT-549 UNISAM**

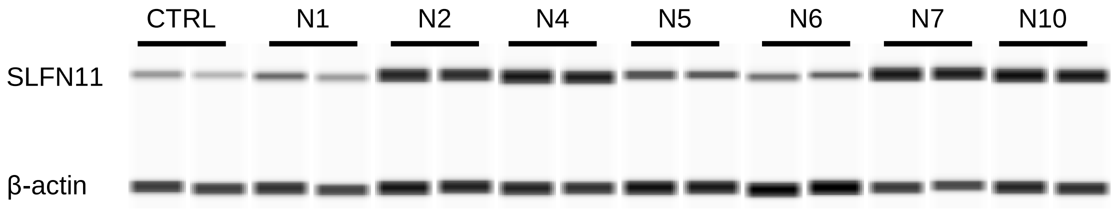


**E** **T47D UNISAM**

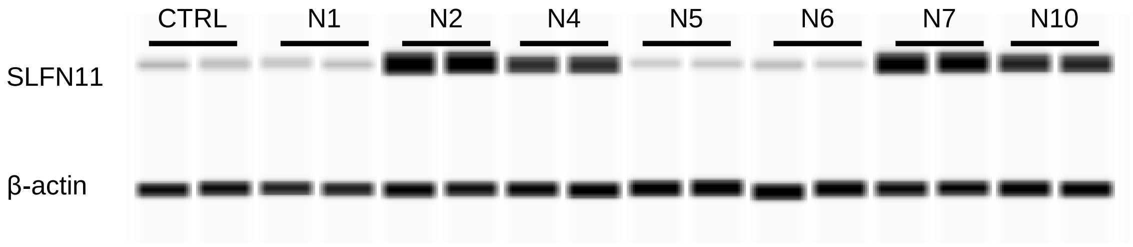


**F** **BT-549 KRAB**

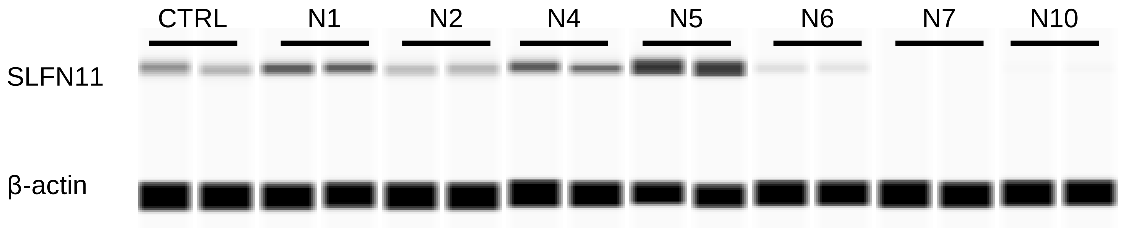


**G** **T47D KRAB**

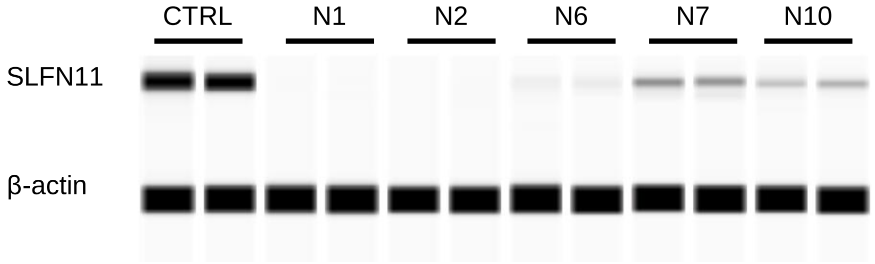

**H**

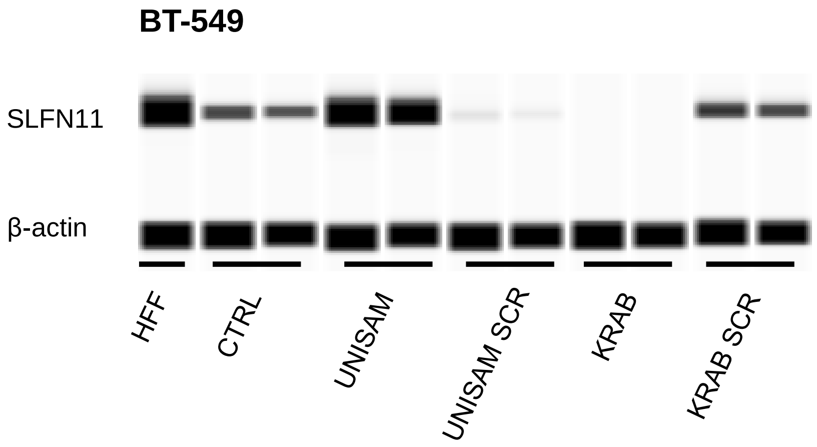
BT-549
SLFN11
β-actin
HFF
CTRL
UNISAM
UNISAM SCR
KRAB
KRAB SCR

**I**

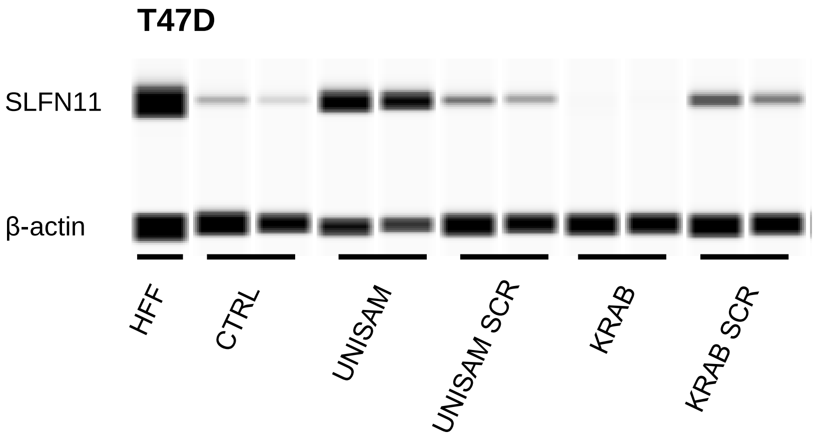
T47D
SLFN11
β-actin
HFF
CTRL
UNISAM
UNISAM SCR
KRAB
KRAB SCR

**J**

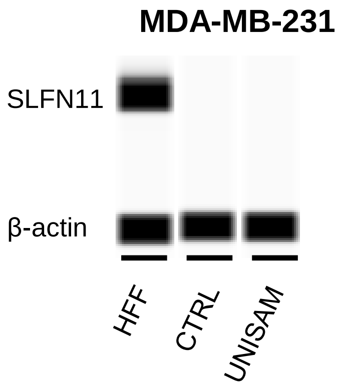
MDA-MB-231
SLFN11
β-actin
HFF
CTRL
UNISAM

### Supplementary Figure 4

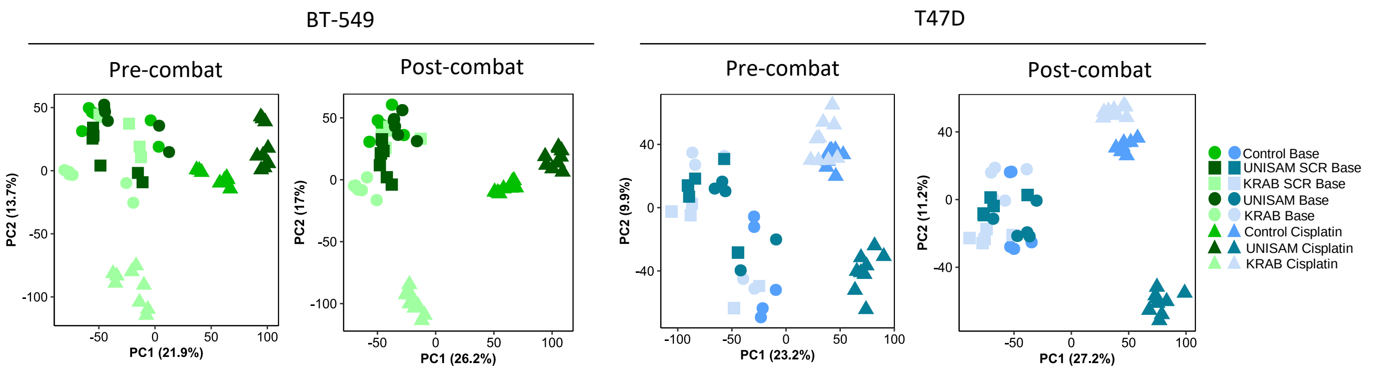

### Supplementary Figure 5

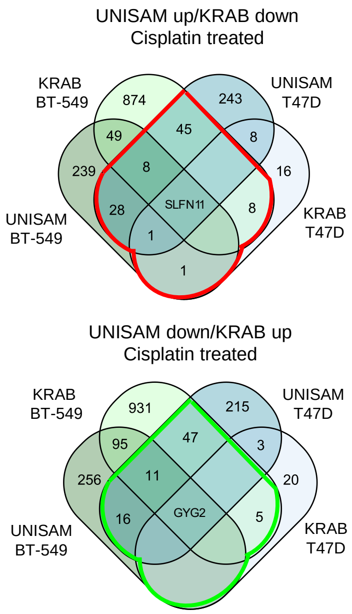


**Cutoff:** FDR < 0.01, LogFC >= 1

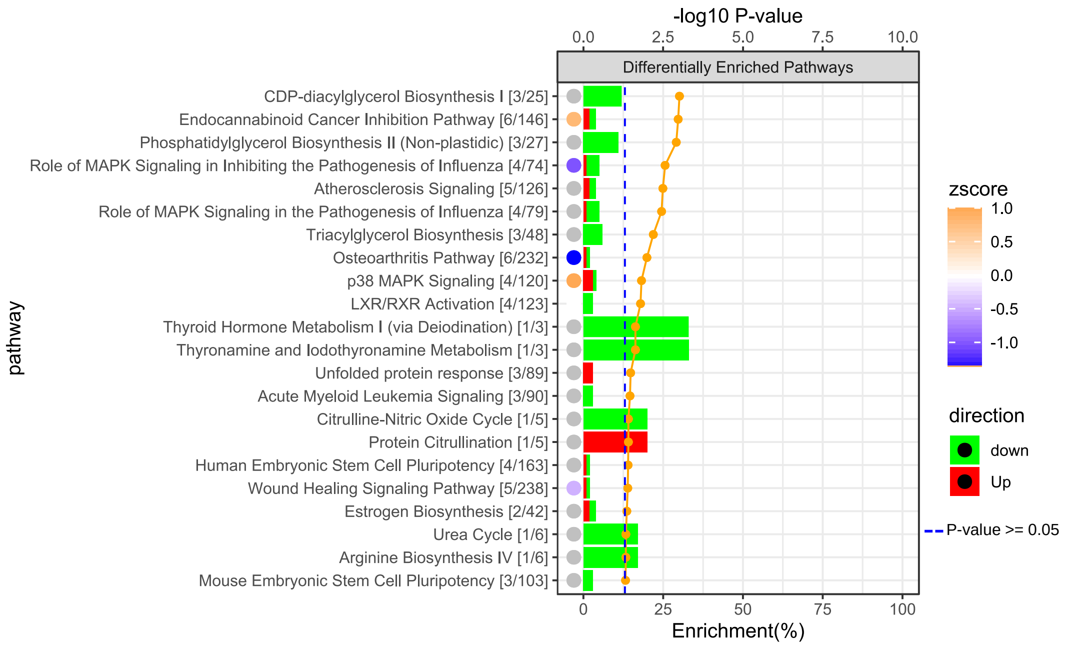
